# Tumors escape immunosurveillance by overexpressing the proteasome activator REGγ

**DOI:** 10.1101/511873

**Authors:** Mathilde Boulpicante, Romain Darrigrand, Alison Pierson, Valerie Salgues, Benoit Gaudineau, Mehdi Khaled, Angela Cattaneo, Angela Bachi, Paolo Cascio, Sébastien Apcher

## Abstract

The success of CD8+ T cell based cancer immunotherapy emphasizes the importance of understanding the mechanisms of generation of MHC-I peptide ligands and possible pathways of tumor cell escape from immunosurveillance. Recently, we showed that peptides generated in the nucleus during the pioneer round of mRNA translation (pioneer translation products, or PTPs) can be a potentially important source of tumor specific peptides, given the presence of aberrant splicing and transcription associated with oncogenesis. Here we show that cancer cells up-regulation of the REGγ proteasome regulator results in increased destruction of PTP-derived peptides in the nucleus thus subverting immunosurveillance. These findings add to understanding of the role of REGγ in antigen processing and identify it as a druggable target for improving the efficacy of cancer immunotherapy.

**Significance:** With the clear success of CD8+ T cell based immunotherapy, it is critical to understand i) how tumor cells generate MHC-I peptide antigens? and ii) the various mechanisms used by cancer cells to evade immunosurveillance. One of them is to up-regulate the REGγ proteasome regulator which results in an increase destruction of MHC-I peptides in the nucleus thus subverting immunosurveillance.

## Introduction

Cellular immune responses against cancer cells expressing non-self epitopes require the activation of CD8+ T cells by professional antigen presenting cells (pAPCs), which take up external peptide material and present it on their major histocompatibility complex class I (MHC-I) molecules as part of a process called cross-presentation (1). The MHC-I direct and cross-presentation pathways are fundamental processes for the detection and elimination of cells that pose a threat to the host. In recent years, it has been suggested that peptides that are directly presented to the MHC-I restricted pathway are not derived from the degradation of full-length proteins but from so-called defective ribosomal products, or DRiPs (2). In addition, some MHC-I-bound peptides are generated by cryptic translation, which refers to polypeptides synthesized in the cell via non-conventional translational mechanisms. These are either peptides encoded by introns, intron/exon junctions, 5’ and 3’ untranslated regions, alternate translational reading frames, or even fusion peptides generated by the proteasome (3–6). These observations have led to a shift in focus towards the notion that the degradation of full-length proteins is not the only critical process for antigen production. More recently, we have shown that the antigen presentation is equivalent whether the peptide is expressed from an intron or from an exon is supported by so-called pioneer translation products (PTPs) (7), which are produced by a translation event distinct from canonical translation prior to mRNA splicing. The PTP model is appealing because it offers an explanation of how the immune system “tolerates” tissue-dependent alternative splicing products. By first examining how direct presentation applies to viral mechanisms of immune evasion, we recently demonstrated that PTPs are also a source of peptides for the exogenous MHC-I pathway. Indeed, PTP-derived peptides are cross-presented by pAPCs in order to specifically activate naïve CD8+T cells (8). Moreover, PTPs are present in exosomes that are engulfed by bone marrow dendritic cells (BMDCs) for cross-presentation. Finally, PTPs purified from tumor cells (tumor-associated PTPs or TA-PTPs) have been used in combination with exosomes as a potent immune cancer vaccine (8).

Polypeptides such as PTPs that enter either the endogenous or the exogenous MHC-I pathway must be processed to fit the MHC-I molecule binding groove to elicit an immune response. A key player of this process is the proteolytic system of the eukaryotic cell, the ubiquitin-proteasome system (9). Central to this system is the proteasome, a multicatalytic complex consisting of a 20S proteolytic core controlled by regulatory complexes that bind to it (10,11). One of these regulatory complexes is the 19S particle, which, along with the 20S proteolytic core, forms the 26S proteasome that degrades ubiquitylated and some non-ubiquitylated proteins in an ATP-dependent manner. Other regulatory complexes, including the REG/PA28 family, have been shown to associate with the 20S or with asymmetric (i.e. single 19S-capped) 26S proteasomes (12,13). The REG family consists of three related subunits, which together form two proteasome regulatory complexes: (i) REGα/β, a heteroheptamer formed by REGα and REGβ subunits, located primarily in the cytoplasm; and (ii) REGγ, a homoheptamer formed by the REGγ subunit, located in the nucleus (14–16). The exact functions and mechanisms of action of REGγ remain elusive, as only a limited number of proteins whose degradation is mediated or controlled by this regulator have been described. Among them are cell cycle regulators, including the cyclin-dependent kinase inhibitors p21 and p16, the oncogene SRC-3 and the tumor suppressor p53 (17–20). This paradigm aligns with the proliferation-promoting and anti-apoptotic properties of REGγ deduced from an analysis of KO mice and the observation that REGγ is overexpressed in many cancers and is often associated with a poor prognosis. Moreover, other observations point to a central role for REGγ in intranuclear dynamics through the regulation of (i) nuclear bodies (including nuclear speckles, Cajal and PML bodies) and (ii) nuclear trafficking of splicing factors (21–23). Evolutionary analyses have shown that REGα and REGβ appeared much later in evolution than REGγ and diverged concomitantly with the emergence of MHC (24). REGγ is important for responses to genotoxic and oxidative stress as well as the impairment of proteasome function, which is for example the case in neurodegenerative diseases (25). Interestingly, it was shown that cellular REGγ is massively recruited to proteasomes after non-toxic treatment with proteasome inhibitors (26). Furthermore, recently, a specific interactor of REGγ has been identified, also called PIP30, which positively modulates the interaction of REGγ to the 20S proteasome complex and alters the selectivity of the REGγ/20S proteasome complex towards different peptides (27). We also recently reported that PTP levels inside the nuclear compartment increased when the proteasomal system was blocked using the highly specific proteasome inhibitor epoxomicin (7).

Although many studies have demonstrated a strong link between REGα/β and antigen production in the immune system (28), no data are currently available suggesting a negative or positive role for REGγ in this process, although REG-knock-out MEF cells and REG-deficient animal models have been produced and challenged to assess specific immune responses (29,30). Here, we show i) an inverse correlation between the expression of REGγ and the presentation of different MHC-I antigenic peptides, ii) REGγ involvement in processing of different PTPs and the negative regulation of CD8+ T cell responses against cancer, iii) a decreased presentation of MHC-I PTP-derived antigens due to the downregulation of REGγ and iv) knockout of REGγ gene causes tumor growth defect *in vivo*. These findings describe a mechanism by which REGγ negatively influences cancer immune responses in an opposite manner compared to the other members of the REG family. These results potentially serve as a starting point for the development of new chemotherapies aiming at decreasing intracellular levels of REGγ thus enhancing the production of tumor-associated antigens and stimulating specific immune responses against cancer.

## Materials and methods

### T cell hybridomas, cell culture and transfection

The SIINFEKL:K^b^-specific (B3Z) and the MBP:K^k^-specific (MBP CD8+) T cell reporter hybridomas were described previously (31,32). Human A375, WM3526 and WM3682 melanoma cell lines, HT29 and T84 colon tumor cell lines, A549 adenocarcinoma lung cells, and MRC5 “normal” lung cells were cultivated in medium recommended by ATCC.

Cells were transfected with different quantities of expression plasmids along with 2 μL of JetPrime according to the manufacturer’s protocol (Ozyme). Each plasmid vector contained a Flag tag.

### Drugs

Cells were treated with different drugs: epoxomicin (Peptides International) was used at 300 nM and cisplastin (Sigma) at 5 and 10 mg/mL.

### T cell assay

Cancer cells were washed twice in 1× PBS and co-cultured either with the SL8-specific B3Z T cell hybridoma or MBP-specific T cell hybridoma for 16–20 h. Then, the cells were centrifuged at 1,200 rpm for 5 min. The cells were washed twice with 1× PBS and lysed for 5 min at room temperature (RT) in the following buffer: 0.2% TritonX-100, 0.5 M K_2_HPO_4_, 0.5 M KH_2_PO_4_. The lysates were centrifuged at 3,000 rpm for 10 min to pellet cell debris. Next, 45 μL of the supernatant was transferred into an optiplate (Packard Bioscience), and a revelation buffer containing 0.01% methylumbelliferyl β-D-galactopyranoside (MUG) was then added. The plate was incubated for 3 h at RT. Human cell lines were transfected with the K^b^ expression vector. Both CD8+ T cell hybridomas express LacZ in response to the activation of T cell receptors specific for the SIINFEKL peptide (Ova-immunodominant peptide) in the context of H2-K^b^ MHC class I molecules or the MBP peptide (myeline basic protein-immunodominant peptide) in the context of H2-K^k^ MHC class I molecules. The activity of β-galactosidase (luminescence) was measured with FLUOstar OPTIMA (BMG LABTECH Gmbh).

### CRISPR/Cas9 transfection and selection

After the transfection of 1 μg of CRISPR plasmid vector, cells were sorted 2 days later (1 cell/well). PCR and Western blotting were performed using the clones; selected clones were sent for sequencing. TOPO TA cloning (Life Technologies) was carried out for selected clones. The CRISPR/Cas9 system was applied to the A375 cell line, and A375 clone number 11 was generated and designated A375c.XI.

### FACS Analysis for H2-K^b^ expression and recovery at the cell surface

To study the kinetics of endogenous surface K^b^ recovery cells were treated with ice-cold citric acid buffer (0.13M citric acid, 0.061M Na_2_HPO_4_, 0.15M NaCl [pH 3]) at 1×10^7^ cells per milliliter for 120s, washed three times with PBS, and resuspended in culture medium. At the indicated time point, an aliquot of cells (generally 1.5×10^6^) was removed and stained with anti-mouse H-2K^b^ PE. All flow cytometry experiments were conducted using the BD LSRII flow cytometer (BD Biosciences) and data are analyzed with the FlowJow software (V10).

### *In vitro* peptide degradation

Reconstitution of REG-20S complexes and the degradation of short fluorogenic substrates, the MP-45 and the KH-52 polypeptides (containing the SIINFEKL epitope in their middle) or the MP-46 and the KH-53 polypeptides (containing the MBP epitope in their middle), were performed according to previously described methods (33–35). Briefly, REGγ- and REGα/β-20S proteasomes were reconstituted by preincubating human 20S constitutive particles (BostonBiochem, USA) with a 6-fold molar excess of REGγ (BostonBiochem, USA) or REGαβ (34) at 37°C for 30 min in 20 mM HEPES, pH 7.6, and 2 mM NaCl and were immediately used for degradation experiments. For kinetic analysis and to generate peptide products for MS/MS studies, all polypeptides (50 μM) were incubated with 20S, REGγ-20S or REGα/β-20S (20 nM) for 8 h at 37°C in 20 mM HEPES, pH 7.6, 2 mM NaCl. To assay the peptides generated during protein degradation, we measured the appearance of new amino groups using fluorescamine as previously described (35). In brief, at the end of the incubation, the peptide products were separated from undegraded polypeptides by ultrafiltration through a membrane with a 3-kDa cutoff (Nanosep, Pall, USA), and these samples were assessed with fluorescamine (Sigma) and used for MS/MS analysis.

### Liquid chromatography–tandem MS (LC–MS/MS) analysis

Five-microliter samples from the 20S -/+ REGγ experiments containing approximately 130 pmol NH_2_/μl were loaded onto StageTipsμC18 (36,37); peptides were eluted in 40 μl 0.1% formic acid. The acetonitrile was allowed to evaporate in a Speed-Vac, and then the samples were resuspended in 6 μl of eluent A (see the composition below) for nLC-MS/MS analysis. Two microliters of each sample was injected as technical replicates into a nLC–ESI–MS/MS quadrupole Orbitrap QExactive-HF mass spectrometer (Thermo Fisher Scientific). Peptide separation was achieved using a linear gradient from 95% solvent A (2% ACN, 0.1% formic acid) to 50% solvent B (80% acetonitrile, 0.1% formic acid) for 23 min and from 60 to 100% solvent B for 2 min at a constant flow rate of 0.25 μl/min on a UHPLC Easy-nLC 1000 (Thermo Scientific) connected to a 25-cm fused-silica emitter with an inner diameter of 75 μm (New Objective, Inc. Woburn, MA, USA), packed in-house with ReproSil-Pur C18-AQ 1.9-μm beads (Dr. Maisch Gmbh, Ammerbuch, Germany) using a high-pressure bomb loader (Proxeon, Odense, Denmark). MS data were acquired using a data-dependent top-15 method for HCD fragmentation. Survey full-scan MS spectra (300–1750 Th) were acquired in the Orbitrap at a resolution of 60,000, an AGC target of 1^e6^, and an IT of 120 ms. For the HCD spectra, the resolution was set to 15,000 at *m*/*z* 200, with an AGC target of 1^e5^, an IT of 120 ms, an NCE of 28% and an isolation width of 3.0 *m*/*z*.

### Data processing and analysis

For quantitative proteomic analysis, raw data were processed with MaxQuant (ver. 1.5.2.8) and searched against a database containing only the sequence of the KH-52 intron SIINFEKL. No enzyme specificity was selected, and there were no differences between I and L. The mass deviation for MS-MS peaks was set at 20 ppm, and the peptide false discovery rate (FDR) was set at 0.01; the minimal length required for a peptide identification was eight amino acids. The list of identified peptides was filtered to eliminate reverse hits. Statistical analyses were performed with Perseus (ver. 1.5.1.6) considering the peptide intensity; normalization based on the Z-score and imputation was applied. Significant peptides were determined with a t-test, Benjamini Hochberg correction and FDR<0.05 (more stringent assignments), or a t-test with a p-value of 0.05 (less stringent assignments). Only significant peptides were used for supervised hierarchical clustering analysis.

### Tumor challenge *in-vivo*

C57Bl/6J female mice were obtained from Harlan Laboratories. NU/NU nude mice were obtained from Charles River. 7 week-old mice were injected subcutaneously into the right flank with 1×10^5^ MCA205 wild type (WT) cells or 1×10^5^ *Cas9-*REG✰ MCA205 clones. Area of the tumor was recorded every 3 to 4 days until ethical limit points are reached. All animal experiments were carried out in compliance with French and European laws and regulations.

## RESULTS

### Inverse correlation between the expression of REGγ and antigen presentation in multiple cancer cell lines

A series of studies have reported the overexpression of REGγ in different cancer types (38–40). To clarify the specific role of REGγ in MHC-I antigen presentation, we selected different cancer cell lines and examined REGγ mRNA and protein expression levels. First, we analyzed REGγ mRNA levels by real-time qRT-PCR in three human melanoma cell lines (A375, WM3526 and WM3682), one human lung cancer cell line (A549) and two colon cancer cell lines (HT29 and T84), and we observed that REGγ was notably upregulated in each cell line compared to the control normal lung cell line MRC5 (Fig. 1A). Next, we measured REGγ protein expression and observed that it was overexpressed in all cancer cell lines compared with MRC5 normal cells, although levels of REGγ overexpression varied between the different cell lines analyzed. Indeed, REGγ was highly overexpressed in melanoma and colon cancer cell lines, expressed at low levels in the lung cancer cell line A549, and demonstrated very low expression in the normal cell line (Fig. 1B). Furthermore, we assessed the cellular distribution of REGγ in these cancer cell lines. Figure 1C shows that endogenous REGγ was localized in the nucleus of all cell lines analyzed independently of its expression levels.

**Figure 1:**
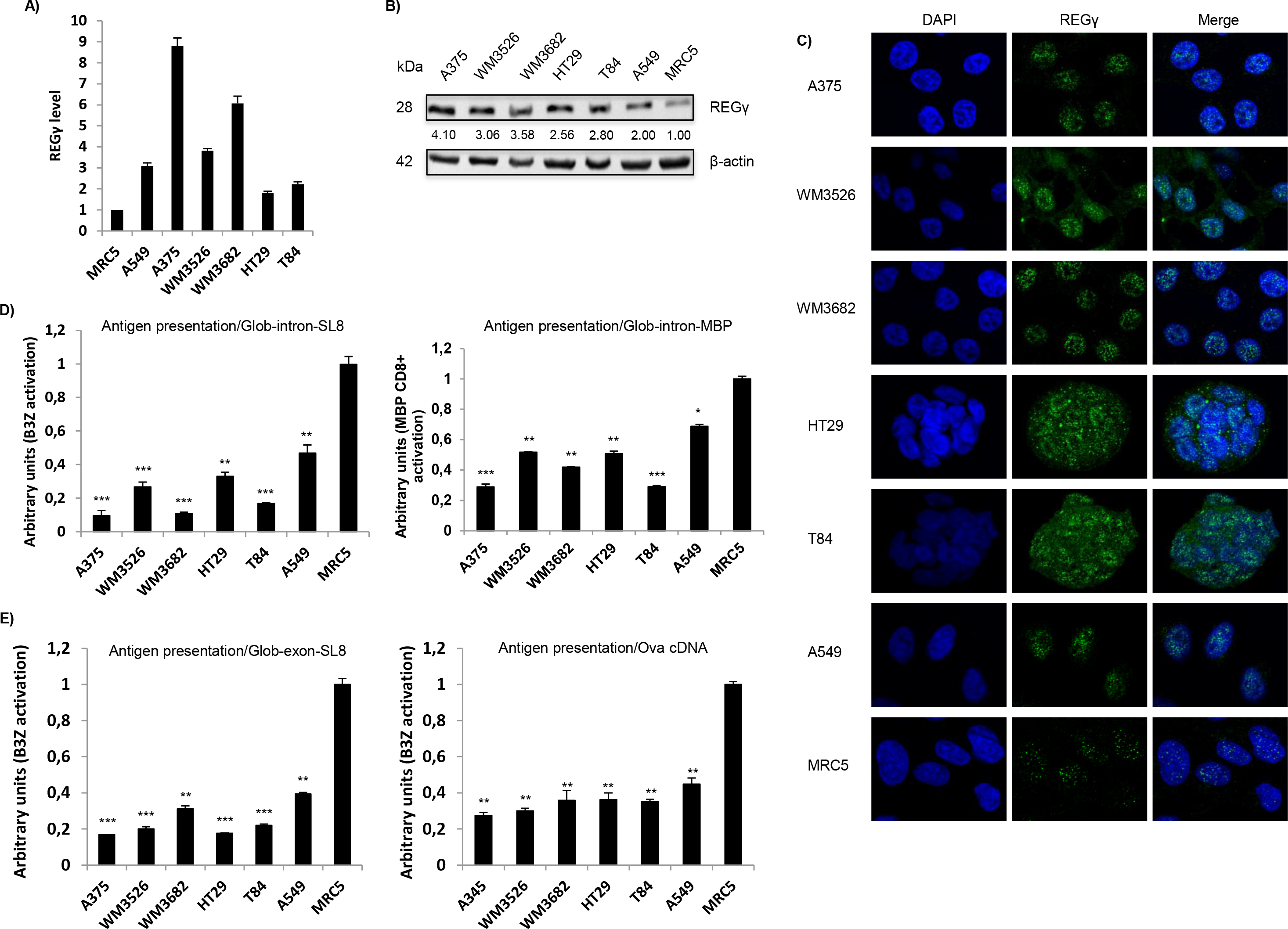
Inverse correlation between the expression of REG γ and antigen presentation in multiple cancer cell lines. **a)** REGγ mRNA levels were analyzed by qPCR in different cell lines and normalized to β-actin mRNA levels. The MRC5 cell line was used as a reference. Experiments were performed in triplicate. Data are expressed as the mean ± SEM from three technical replicates. **b)** Western blot analysis and quantification (relative to the housekeeping protein β-actin) of REGγ expression. Protein levels are indicated below each gel. **c)**REGγ protein localization was determined by immunofluorescence in several tumor cell lines and in the MRC5 normal cell line. REG was stained with Alexa Fluor 488, and nuclei were counterstained with DAPI. Cells were analyzed by confocal microscopy. As expected, the regulator REGγ was localized in the nucleus in all tested cell lines. **d, e)** All cell types were transfected *in vitro* with β-Glob-intron-SL8 (**d, left panel**) or β-Glob-intron-MBP(79-87) (**d, right panel**), β-Glob-exon-SL8 (**e, left panel**) and OVA cDNA (**e**, **right panel**) expression constructs. The cells were incubated with the SL8-specific CD8+ T cell hybridoma (B3Z) for 16 h. The data show the average of at least three independent experiments ± SD minus the values from mock-transfected cells. Free SL8 peptides were added to the cells to ensure that the T cell assays were performed under non-saturation conditions and that the expression of MHC-I molecules was not affected.

To evaluate whether the observed differences in REGγ expressions in cancer and normal cell lines contribute to changes of antigen production and presentation, we examined the presentation of PTP-derived-antigens at the cell surface of these cell lines. To achieve this goal, we expressed the mouse MHC-I K^b^ molecule and Glob-intron-SL8 constructs in these human cell lines, which enabled us to examine the production of specific SL8-comprising-PTPs as described previously (6,7). Using a SIINFEKL:K^b^ (B3Z) T cell hybridoma (31) that specifically detects the SL8 epitope presented on K^b^ molecules, we first found that the antigen presentation was lower in all cancer cell lines tested than in the normal MRC5 cell line (Fig. 1D, left panel). Interestingly, we observed a complete inverse correlation between PTP-derived-SL8 presentation and REGγ expression in all cell lines tested. This result suggested that REGγ overexpression negatively affected PTP-derived antigen presentation (compare Figs. 1B and 1D). Crucially, this result was not merely due to differences in the cellular level of the β-globin construct, which in fact were higher in tumor cell lines overexpressing REGγ (Supplementary Fig. 1A). Moreover, FACS analysis showed that overall H2-K^b^ expression differed among the cancer cell lines tested, but it did not correlate with the expression of SL8 as determined with the B3Z assay. In fact, we observed that MHC-I molecules were more abundant at the cell surface of tumor cell lines overexpressing REGγ and exhibiting a decrease in PTP-dependent antigen presentation (Supplementary Fig. 1B).

In order to rule out the possibility that the negative role of REGγ on PTP dependent antigen presentation could be restricted to the SL8 epitope or the K^b^ molecule, we determined whether also the presentation of the MBP(79-87) epitope, which is derived from the Myelin Basic Protein (MBP) and is presented on K^k^ molecules was affected by the overexpression of REGγ in different cancer cell lines. Using the specific MBP CD8+ T cell hybridoma (32) we could obtain results similar to those previously observed with the PTP-derived-SL8, that is to say a complete inverse correlation between PTP-derived-MBP presentation and REGγ expression in all cancer cell lines tested (Fig. 1D, right panel). All together, these results indicate: i) a negative role of REGγ in the MHC-I antigen presentation pathway since in all cell lines overexpressing it the β-globin protein and the MHC-I molecules were more abundant while the antigen presentation was reduced and ii) that the inverse correlation between REGγ expression and antigen presentation is not restricted to a specific epitope or MHC-I molecule.

We then wanted to determine whether the previously observed differences in SL8 antigen presentation were restricted to intron-derived tumor-associated antigens or if they were also observed with an exon-derived tumor-associated antigen. To this purpose, we analysed cells expressing mouse MHC-I K^b^ molecule and Glob-exon-SL8 construct or ovalbumin cDNA, in which the SL8 epitope is found in its correct setting. As shown in Figure 1E, antigen presentation showed an inverse correlation with REGγ expression in all cell lines tested, independently of the position of the antigenic epitope in the exon sequence (left panel) or in cDNA constructs (right panel).

Furthermore, we expressed in the different human cell lines the mouse MHC-I H2-K^b^ molecule and the SL8-minigene construct, which contains only the presented 8-amino-acid sequence of SL8, and observed that differences in REGγ expression did not correlate with SL8 antigen presentation (Supplementary Fig. 1C). Hence, the effects of REGγ on MHC-I antigen presentation only occurred with longer polypeptides containing the presented MHC-I antigenic peptides, indicating that REGγ directly acts on epitopes processing and not on subsequent steps of class-I presentation.

All together these results clearly support the hypothesis that the proteasome regulator REGγ is involved in the inhibition of MHC-I presentation of antigens present in exon as well as in intron-derived PTPs, a mechanism that may allow tumor cells to avoid immune responses against conventional and unconventional epitopes.

### Exogenous overexpression of REGγ decreases MHC-I antigen presentation

In the rest of our study we decide to use the Glob-intron SL8 construct since, due to its nuclear localization it is the most appropriate tool to investigate the role of a nuclear proteasomal regulator on PTPs dependent antigen presentation. To study the specific effects of REGγ on PTP-dependent MHC-I antigen presentation, MRC5 lung fibroblast cells and A549 lung cancer cells, which express REGγ at very low levels, (see Figs. 1A and 1B), were therefore co-transfected with the mouse MHC-I K^b^ molecule, the Glob-intron-SL8 and an exogenous Flag-REGγ construct. Using, Western blotting (Fig. 2A), immunofluorescence (Fig. 2B) or qRT-PCR (Supplementary Figure 2), we monitored the expression of the exogenous REGγ construct and compared it to the expression of endogenous REGγ. Notably, the overexpressed exogenous REGγ showed the same expression pattern and localization as the endogenous REGγ. Next, we examined the antigen presentation of the SL8 epitope in these cell lines. The introduction of increasing amounts of REGγ in both A549 and MRC5 cell lines, which express fixed amounts of the Glob-intron-SL8 and MHC-I K^b^ molecules, resulted in a dose-dependent decrease in the activation of the B3Z hybridoma (Fig. 2C). This observation supports the hypothesis that the overexpression of REGγ has a clear-cut negative effect on PTP-dependent antigen presentation.

**Figure 2:**
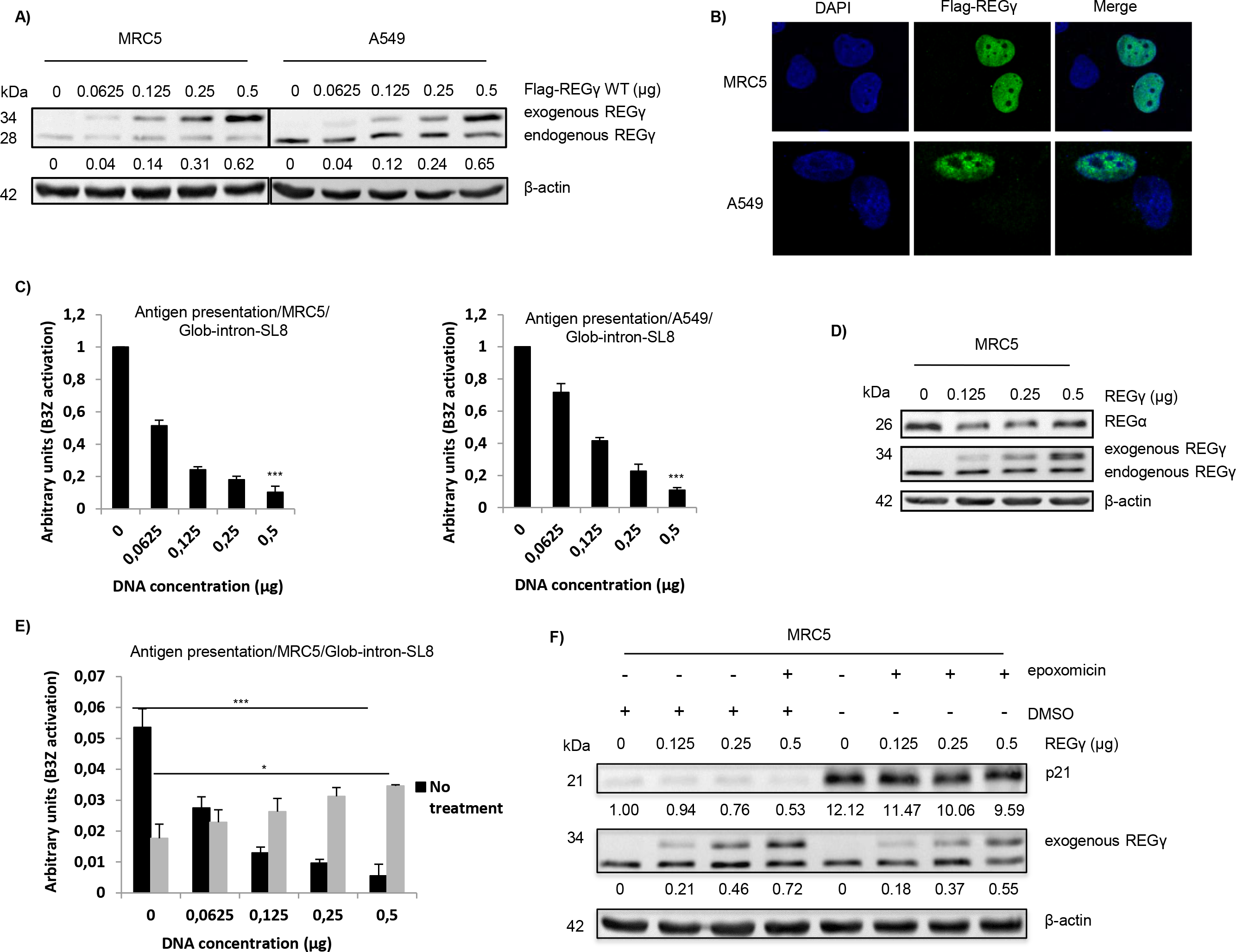
Exogenous REGγ overexpression decreases antigen presentation. **a)** MRC5 and A549 were transfected with a construct expressing REGγ WT from 0.0625 to 0.5 μg or a corresponding empty construct for 48 h. REGγ protein levels were examined by Western blotting using β-actin as a loading control. Protein levels are indicated below each gel. **b)** MRC5 and A549 were transfected with a construct expressing Flag-REGγ WT (0.25 μg). The Flag tag was stained with Alexa Fluor 488, and the nuclei were stained with DAPI. Cells were analyzed by confocal microscopy. As expected, exogenous transfected REGγ WT localized in the nuclei. **c)** MRC5 cells (left panel) and A549 cells (right panel) were transfected with the Glob-intron-SL8 construct (0.5 μg). The cells were incubated with the SL8-specific CD8+ T cell hybridoma (B3Z) for 16 h. The data show the average of at least three independent experiments ± SD minus the values from mock-transfected cells. **d)** MRC5 cells were transfected with a REGγ WT construct for 48 h. The increase in REGγ protein expression had no effect on the expression of REGα protein. **e)** MRC5 cells were transfected with increasing amounts (from 0 to 0.5 μg) of a Glob-intron-SL8 construct for 48 h. At 36 h post-transfection, the cells were μ then treated overnight with epoxomicin (300 nM). Next, the cells were incubated with the B3Z T cell hybridoma for 16 h. The data show the average of at least three independent experiments ± SD. *** p < 0.001, * p < 0.05 (unpaired t-test). **f)** Western blot analysis and quantification (relative to the housekeeping protein β-actin) of p21 expression in MRC5 cells treated overnight with epoxomicin at 300 nM. Protein levels are indicated below each gel. As expected, p21 protein levels increased after 12 h of epoxomicin treatment, even though the treated cells overexpressed the exogenous regulator REGγ.

As mentioned earlier, REGγ is not the sole regulator of the proteasomal pathway; 20S proteasome also binds to the REGα/β regulator, which has been shown to be implicated in the accurate processing of MHC-I antigenic epitopes (14). To investigate more in detail at which exact level of the proteasomal pathway REGγ contributes to PTP-dependent antigen presentation regulation, we determined whether the overexpression of REGγ influenced the expression of endogenous REGα. Figure 2D shows that increasing the amount of exogenous REGγ did not affect the expression of REGα. This result therefore demonstrates that the decrease in PTP-dependent antigen presentation causes by REGg in different cancer cell lines is not due to a decrease in the expression of the regulator REGα but more likely it is the result of a direct effect on the 20S proteasome. In fact, the presentation of antigenic peptides depends on proteolytic processing by the 20S proteasome complex. To ensure that differences in the production of SL8 peptides by the proteasome are unambiguously attributable to the overexpression of REGγ we treated MRC5 cells expressing increasing amounts of REGγ and a fixed amount of Glob-intron-SL8 and MHC-I K^b^ molecules with the 20S proteasome-specific inhibitor epoxomicin and examined SL8 presentation. Interestingly, while non-treated cells displayed a decrease in antigen presentation with REGγ overexpression, a partial increase in MHC-I antigen presentation was observed when cells were treated with epoxomicin (Fig. 2E). This observation supports the idea that REGγ does not exert unspecific processing effects on MHC class I antigen presentation in the overexpressing cell lines but instead confers a real and specific effect on the 20S core proteasome. Moreover, as a control for the real efficacy of the proteasomal inhibitor under the condition used in the treated cell lines, we examined p21 protein levels, which has been reported to be regulated specifically by the REGγ-20S proteasome complex in an ubiquitin and ATP-independent process (18). p21 protein levels increased following 18 h of epoxomicin treatment, even though the treated cells overexpressed the exogenous REGγ regulator (Fig. 2F).

Therefore, all these results confirm that the overexpression of REGγ does not result in (i) a decrease in MHC-I expression or export, or (ii) a decrease in the expression of the other members of the REG family, but does directly affect proteasomal proteolytic activities.

### Knockdown and knock-out of the regulator REGγ promote antigen presentation

Considering our finding showing that endogenous overexpression of the REGγ regulator negatively affects MHC-I antigen presentation in cancer cells, we speculated whether REGγ knockdown might restore PTP-dependent antigen presentation. To test this hypothesis, we silenced REGγ by transient siRNA treatment in human melanoma and colon cancer cell lines. As expected, REGγ knockdown (Fig. 3A) enhanced p21 protein levels in all cancer cell lines compared to scrambled siRNA (Fig. 3B). More importantly in terms of PTP-dependent antigen presentation, we observed a close correlation between REGγ knockdown and the increase of the SL8 epitope at the cell surface (Fig. 3C).

**Figure 3:**
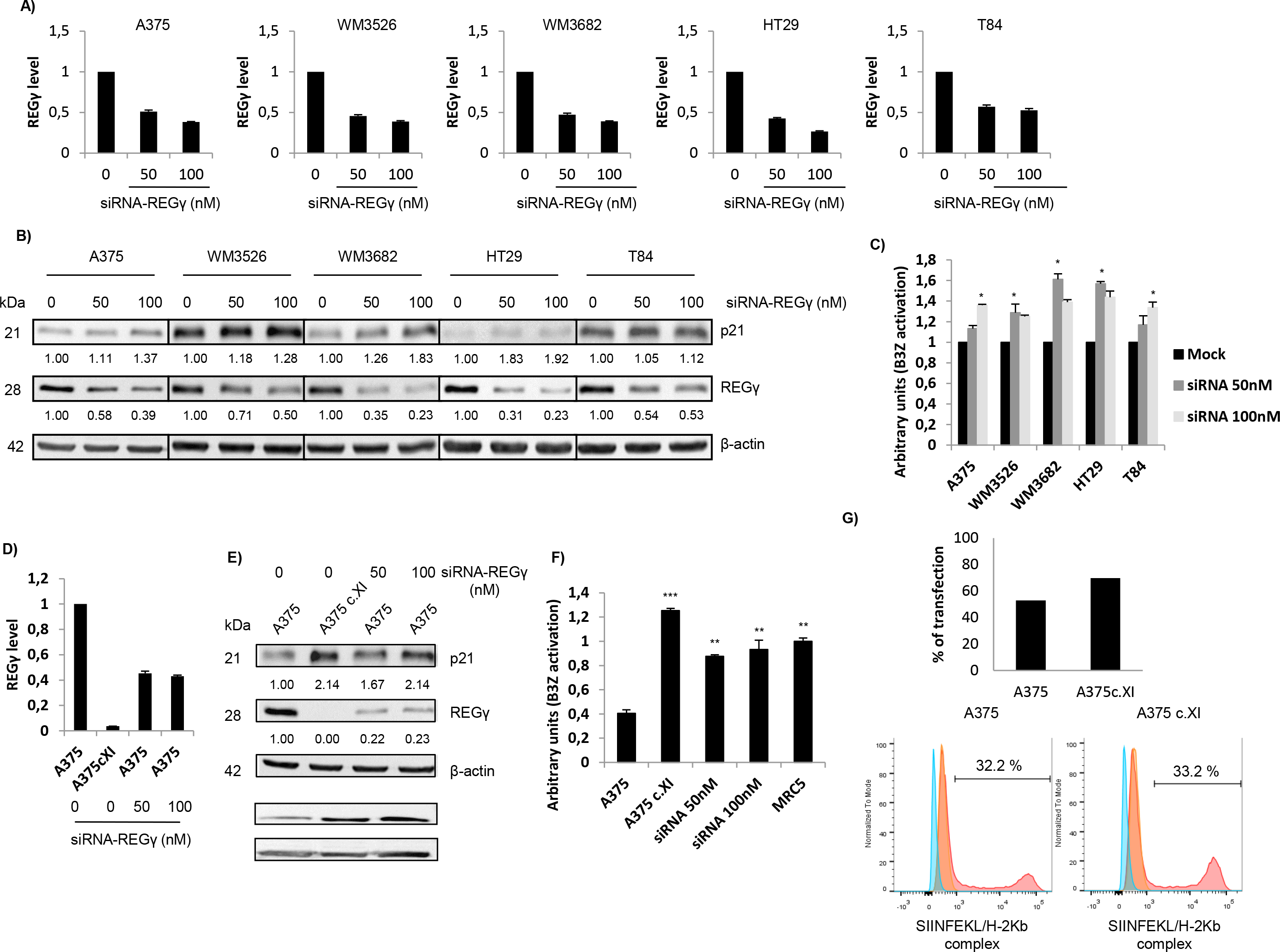
Knockdown and knock-out of the expression of the regulator REGγ promotes antigen presentation. The A375, WM3526, WM3682, HT29 and T84 cell lines were transfected with siRNA specific for REGγ (50 and 100 nM) or siRNA negative control (0) for 48 h and then analyzed by RT-qPCR (**a**), Western blotting (**b**) and a T cell assay (**c**). **a)** Experiments were performed in triplicate. Data are expressed as the mean ± SEM of three technical replicates. **b)** Western blotting was performed to analyze and quantify REGγ and p21 expression, and β-actin was used as a reference protein. The relative protein level is indicated below each gel. **c)** Cell lines were transfected with the Glob-intron-SL8 construct (0.5 μg) and treated with different concentrations of siRNA specific for REGγ (0, 50 and 100 nM). After 48 h, the different cell lines were incubated with the B3Z T cell hybridoma for 16 h. The data show the average of at least three independent experiments ± SD minus the values from mock-transfected cells. **d)** qPCR analysis of A375 cells transfected with siRNA specific for REGγ (50 and 100 nM) was performed. The inhibition of REGγ mRNA levels was quantified and normalized to β-actin mRNA levels. Experiments were performed in triplicate. Data are β expressed as the mean ± SEM from three technical replicates. **e)** Western blot analysis and quantification (relative to the housekeeping protein β-actin) of REGγ and p21 expression. Protein levels are indicated below each gel. **f)** The A375cXI and A375 cell lines were both transfected with the Glob-intron-SL8 construct (0.5 μg) for 48 h, and REGγ-siRNA (50 or 100 nM) was only added to A375 cells. The cells were incubated with the B3Z T cell hybridoma for 16 h. The data show the average of at least three independent experiments ± SD minus the values from mock-transfected cells. *** p < 0.001, ** p < 0.01, * p < 0.05 (unpaired t-test). **g)** A375 and A375cXI cells were transfected with the Glob-intron-SL8 construct vector for 48 h. The transfection efficacy was analyzed by FACS (top panel). Human tumor cell lines were transfected with a construct expressing the H-2K^b^ gene and stimulated with extracellular SIINFEKL synthetic peptide for 15 min. The transfection efficacy of the MHC class I K^b^ molecules was analyzed by FACS (bottom panels). Controls were stained with mouse IgG1 K isotype control APC. Data are presented as percentages.

The expression of REGγ is controlled by several mechanisms inside the cell. miRNA-7, which has been reported as a tumor suppressor (41,42) and is downregulated (43,44) in several human cancers, also negatively controls the expression of the regulator REGγ in lung cancer (45). The A375 melanoma cancer cell line, which in our hands expressed the highest levels of REGγ, was therefore transfected with increasing amounts of miRNA-7. Of great interest, we observed that miRNA-7 inhibited REGγ expression (Supplementary Fig. 3A) and led to an increase in PTP-SL8-dependent antigen presentation (Supplementary Fig. 3B) (41,42)(41,42)(41,42)(41,42)(41,42)(40,41)(38,39).

In addition, treatment of cancer cells with the chemotherapeutic molecule cisplatin has been shown to induce a decrease in mRNA and protein levels of REGγ (46) (and Supplementary Fig. 3C). Accordingly, treatment of A375 cells with cisplatin restores the expression of PTP-dependent antigens at the cell surface (Supplementary Fig. 3D). All together, these results support the idea that REGγ downregulation restores PTP-derived antigen presentation and indicate a possible chemotherapeutic approach to achieve this goal. To clarify ore in detail the functional importance of REGγ in cancer immune escape, we used the CRISPR/Cas9 system to create a melanoma *Cas9*-A375 REGγ-knock-out cell line (A375cXI). The heterozygous knock-out A375cXI was confirmed by Sanger sequencing, qRT-PCR and Western blot analysis. The A375cXI cell line did not express REGγ mRNA (Fig. 3D) or its corresponding protein (Fig. 3E). As a control for the loss of REGγ expression, the p21 protein accumulates in this cell line, as in cells that were knocked down with specific REGγ siRNAs compare to the parental A375 WT line (Fig. 3E). Crucially, when analyzed for antigen presentation and processing, A375cXI cells proved able to strongly activate the PTP-dependent immune response in contrast to A375 WT (Fig.3F). Moreover, the level of antigen presentation in these cell lines was even higher than the control MRC5 line, which weakly expresses REGγ (Fig. 3F). Differences in REGγ levels are likely to account for this result, since both the tested cell lines (A375cXI and A375 WT) expressed equal amounts of exogenous Glob-intron-SL8 protein (Fig. 3G, top panel) as well as equal amounts of the mouse MHC class I molecule at their cell surface (Fig. 3G, bottom panel).

All together these results clearly confirm that there is an inverse correlation between the expression of REGγ and PTP-dependent antigen presentation in cancer cell lines. This effect is likely due to a close control of the proteasomal degradative pathway and not to an overall effect of REGγ on MHC-I molecules expression and export to the cell surface.

### REGγ regulates the nuclear proteasomal pathway

In our previous studies, we demonstrated that PTPs are produced by a translation event that is distinct from canonical translation and occurs prior to mRNA splicing, supporting the idea that PTPs are generated by a nuclear translation event. Furthermore, we have also reported that treatment with the proteasome inhibitor epoxomicin increases the amount of PTPs within the nuclear compartment, indicating that the nuclear proteasome may be involved in the processing and/or degradation of PTP-antigenic epitopes (7). Based on these studies and all our results so far, we next sought to understand how REGγ contributes to the specific inhibition of PTP-dependent cancer immune responses and in which cell compartment this inhibition occurs. Antigenic epitopes derived from PTP processing are generated either by a cytosolic or nuclear proteasomal complex, in which the REG family plays an essential role. As proteasomal regulator, REGγ has to bind to the 20S proteasome to control protein or polypeptide degradation and this interaction should occur in the nucleus. A REGγ gene sequence mutation that leads to the replacement of Asn 151 by Tyr (N151Y) has been reported to impair the ability of the regulator to activate the trypsin-like activity of the 20S proteasome (47). A375cXI cells were transfected with increasing amounts of Flag-REGγ WT or mutated Flag-REGγN151Y. First, we showed by western-blotting that compared to the WT REGγ the mutated regulator was no longer able to degrade the p21 protein likely as a direct consequence of its inability to activate the trypsin-like activity of the 20S proteasome. Moreover, the expression of the other members of the REG family was not impacted under all the experimental conditions used (Fig. 4A). More importantly, Flag-REGγN151Y inhibited SL8 antigen production and presentation at the cell surface from Glob-intron-SL8 less efficiently than Flag-REGγ WT (Fig. 4B). This inhibition of SL8 presentation again was not attributable to a decrease in expression of the other REG family members shown to be involved in antigen presentation. Two explanations are therefore consistent with this result: i) exogenous mutated Flag-REGγN151Y may no longer be localized in the nucleus; or ii) exogenous mutated Flag-REGγN151Y may have lost its capacity to bind to the 20S proteasome and, therefore, contrary to Flag-REGγ WT it is no longer able to inhibit proteasome activity. To discriminate between these two alternative hypotheses, we first examined the distribution of the mutated Flag-REGγN151Y as well as its capacity to bind to the 20S proteasome *in cellulo*. As shown in Figure 4C, the mutated and WT exogenous REGγ regulators are both localized in the nucleus. Since we did not observe any differences in the localization that could explain the differences in the production of MHC class I epitopes between the cell lines expressing the mutated or the WT regulators, we determined whether mutated Flag-REGγN151Y retained the capacity to bind to the nuclear 20S proteasome *in cellulo*. To achieve this goal, we used proximal ligation assay (PLA)-labeled secondary antibodies, which allow the detection of two primary antibodies in close proximity. Using a combination of PLA-labeled anti-Flag and anti-proteasomal α4 antibodies, we observed a specific signal in the nuclear compartments of cells expressing the mutated and WT REGγ constructs, revealing the co-localization of both form of REGγ with the 20S proteasome (Fig. 4D). Furthermore, as a control, anti-Flag alone under these conditions produced no PLA reaction, and no positive signals were detected in non-transfected cells (Fig. 4D, top panel). We confirmed further the interaction between WT Flag-REGγ and mutated Flag-REGγN151Y with the 20S core proteasome by coimmunoprecipitation using an antibody directed against the REGγ regulator (Fig. 4E). Localization and capacity to bind to the 20S core were equivalent between exogenous Flag-REGγ WT and mutated Flag-REGγN151Y, again demonstrating that the REGγ-20S proteasome complex is involved in the processing/degradation of PTP-derived antigenic peptides.

**Figure 4:**
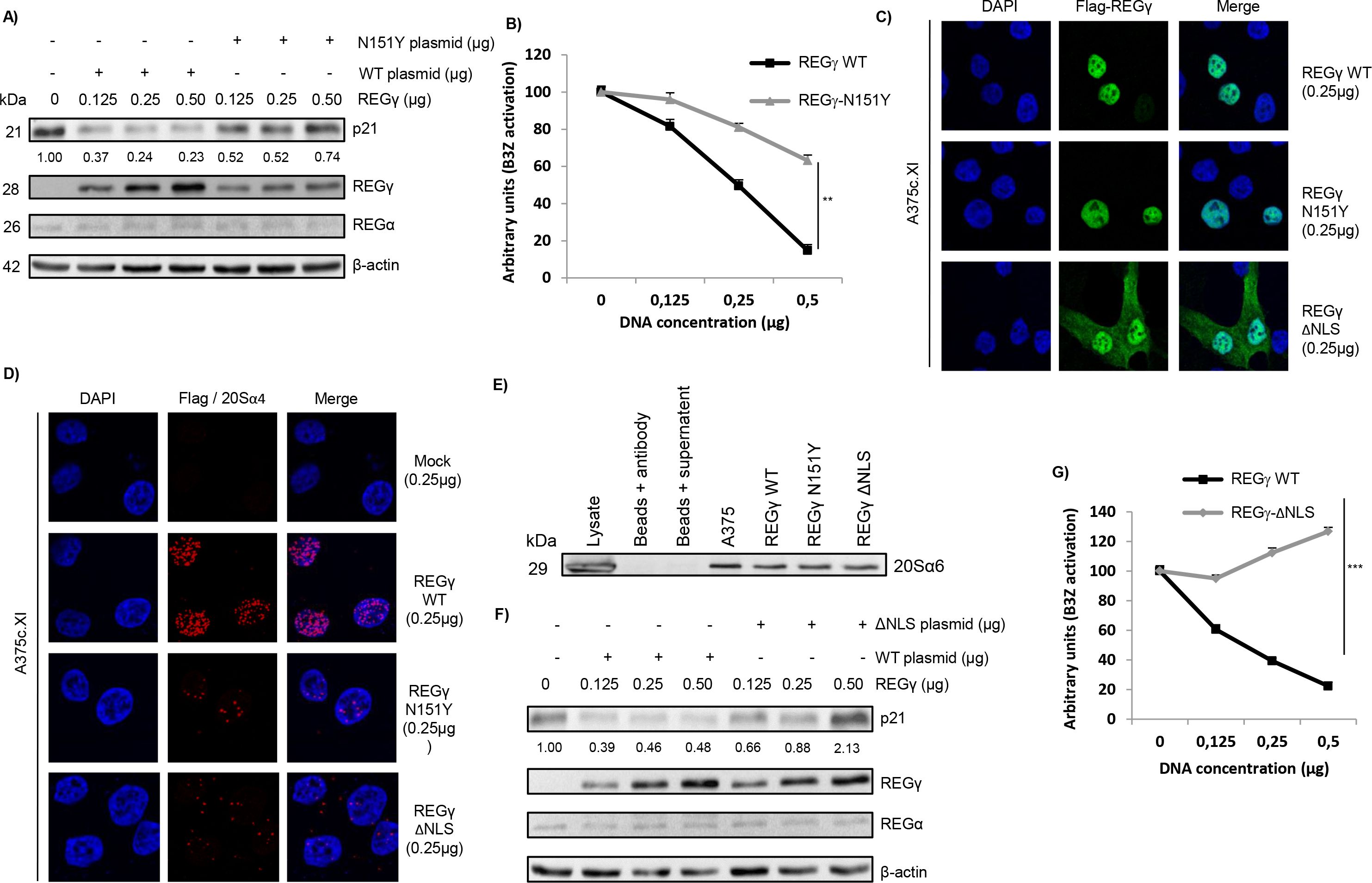
REGγ regulates the nuclear proteasomal pathway. **a)** Western blotting was performed to analyze and quantify REGγ, REGα and p21 protein expression in A375cXI cells (relative to the housekeeping protein β-actin). The cells were transfected with a construct expressing REGγ WT or the mutated REGγN151Y at different concentrations or with a corresponding empty construct for 48 h. Protein levels are indicated below each gel. **b)** Cells were transfected with the Glob-intron-SL8 construct (0.5 μg) and then were incubated with the B3Z T cell hybridoma for 16 h. Data are the μ average of at least three independent experiments ± SD minus the values from mock-transfected cells. A375cXI cells were transfected with a construct expressing a Flag-tagged REGγ WT or a Flag-tagged REGγ N151Y. The Flag tag was stained with Alexa Fluor 488, and the nuclei were stained with DAPI. Cells were analyzed by confocal microscopy. As expected, the exogenous REGγ WT and REGγN151Y proteins were localized in the nuclei. **d)** The interactions of the 20S proteasome with the exogenous REGγ WT and REGγN151Y constructs were analyzed using DuoLink. PLA was performed using an antibody against the Flag tag and α4 subunit - of the 20S proteasome. Staining was analyzed by confocal microscopy. **e)** Immunoprecipitation was performed using the REGγ antibody, and Western blotting was carried out using an antibody against the 20S_α6 subunit. This experiment confirmed the interaction of the 20S proteasome and exogenous REGγ WT or REGγN151Y. **f)** Western blot analysis and quantification (relative to the housekeeping protein β-actin) of REGγ, REGγ and p21 protein levels in A375c.XI cells transfected with a construct expressing REGγ WT or REGγΔNLS at different concentrations or with a corresponding empty construct for 48 h. Protein levels are indicated below each gel. **g)** Cells were transfected with Glob-intron-SL8 construct (0.5 μg). The cells were then incubated with the B3Z T cell hybridoma for 16 h. The data show the average of at least three independent experiments ± SD minus the values from mock-transfected cells.

To further define the compartment hosting REGγ-dependent PTP degradation, we generated an exogenous Flag-REGγ construct that did not contain a nuclear localization signal (NLS). Unlike Flag-REGγ WT, Flag-REGγΔNLS was expressed throughout the cell and was not confined to the nucleus (Fig. 4C). Interestingly, the expression of Flag-REGγΔNLS was not confined to the cytoplasm, suggesting that REGγ contains more than one nuclear localization signal. The combination of PLA-labeled anti-Flag and anti-proteasomal α4 antibodies produced a specific signal in the cytoplasmic compartments of cells expressing the Flag-REGγΔNLS and Flag-REGγ WT constructs. We also observed staining corresponding to an interaction between Flag-REGγΔNLS and the 20S core in the nucleus. This result indicates that Flag-REGγΔNLS, which was localized throughout the cell, retained the capacity to bind to the 20S proteasome in both the cytoplasm and the nucleus (Fig. 4D). We also confirmed the interaction between Flag-REGγ WT and Flag-REGγΔNLS with the 20S core proteasome by coimmunoprecipitation using an antibody directed against the REGγ regulator (Fig. 4E). Moreover, A375ΔREGγ cells transfected with Flag-REGγΔNLS slightly prevented nuclear 20S proteasome degradation of the p21 protein in a REGγ-dependent manner compared to cells transfected with Flag-REGγ WT (Fig. 4F). This result suggests that the low levels of Flag-REGγΔNLS in the nucleus retained the capacity to partially activate the nuclear 20S proteasome and, furthermore, demonstrate that the degradation of the p21 protein is a nuclear and not a cytoplasmic event (Fig. 4F). In parallel, when we examined the effects of the partial relocalization of Flag-REGγΔNLS in the cytoplasm on the antigenic presentation pathway, we observed an increase in SL8-PTP-dependent antigen presentation compared to the loss of antigen presentation in cells expressing exogenous nuclear Flag-REGγ WT (Fig. 4G).

All of these results hence demonstrate that REGγ is involved in PTP-dependent antigen presentation by binding to the 20S proteasome in the nucleus.

### REGγ promotes the degradation of MHC class I PTP-derived antigenic epitopes

The above results raised at least two alternative hypothesis regarding the role of REGγ in the PTP-dependent proteasomal degradation pathway: i) REGγ may inhibit 20S proteasomal peptidase activities are responsible for producing antigenic peptides with the correct characteristics (i.e. length and anchor residues) to serve in MHC-I pathway, with a consequent decrease in production of PTP-dependent epitopes; or ii) REGγ may stimulate specific 20S proteasomal activities that cause further degradation of the correct size MHC-I antigenic epitopes. Recently, we showed that inhibition of the proteasomal pathway with epoxomicin leads to increased amounts of unprocessed PTPs in cells and promotes the cross-presentation of longer polypeptides compared with untreated cells (8). To test the first hypothesis, we transfected different cancer cell lines, normal MRC5 fibroblasts and the A375ΔREGγ cell line with the Glob-intron-SL8 construct and incubated them with mouse BMDCs for 24 h. We observed a reduction in B3Z activation when the tumor cell lines were incubated with BMDCs compared to the activation levels observed with A375ΔREGγ and MRC5 cells (Fig. 5A). The decrease in cross-presentation in the different cell lines overexpressing REGγ suggested that REGγ did not inhibit PTP processing, leading to an accumulation of unprocessed polypeptides, which would have been a source of material for the exogenous MHC-I pathway.

**Figure 5:**
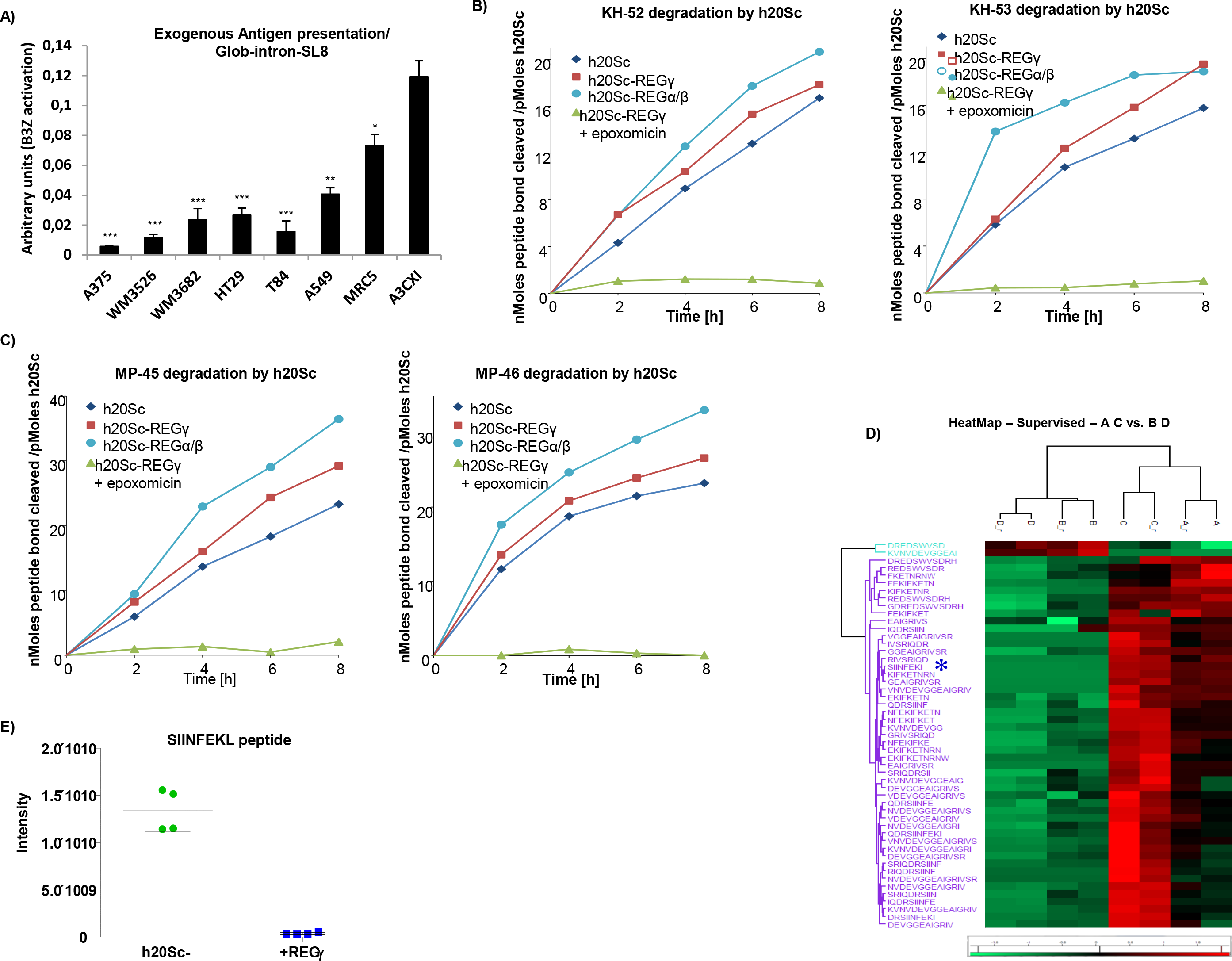
REGγ promotes the degradation of MHC-I PTP-derived antigenic epitopes. **a)** A375, WM3526, WM3682, HT29, T84, A549, MRC5 and A375cXI cell lines expressing the Glob-intron-SL8 construct (0.5 μg) were cultured with BMDCs for 24 h. The BMDCs were then co-cultured with the SL8-specific CD8+ T cell hybridoma (B3Z) for 16 h, and T cell activation was estimated by measuring β-galactosidase levels. The data show the average of at least three independent experiments ± SD minus the values from mock-transfected cells. *** p < 0.001, ** p < 0.01, * p < 0.05 (unpaired t-test). Hydrolysis rates of KH-52 and KH-53 (**b**) and MP-45 and MP-46 (**c**) peptides. Precursor substrates were incubated with human 20S constitutive proteasomes (h20Sc) alone or activated by REGγ or REGα/β, and the amino groups released were measured with fluorescamine at the indicated time points. Data are representative of three independent experiments. **d)** Heatmap comparison of the abundance of significant peptides generated from proteasomal degradation of KH-52 precursor peptide in the h20Sc (biological and technical replicates of samples designated A and C, respectively) and REGγ-h20Sc (biological and technical replicates of samples designated B and D, respectively) samples. Differences and similarities in peptide intensities (normalized to the Z-score) are shown; green indicates decreased levels, and red indicates increased levels. Data were obtained from supervised hierarchical clustering analysis by applying a t-test and a p-value of 0.05. **e)** Box plot of SIINFKEL peptide intensity calculated by MaxQuant in the h20Sc (in green) and REGγ-h20Sc (in blue) samples.

To confirm that REGγ did not inhibit the capacity of 20S proteasome to hydrolyze long polypeptides containing epitopes, we assessed the effects of this regulator on the 20S core particle cleaving properties by using an *in vitro* degradation assay. To achieve this goal, a 45-mer (MP-45), and a 52-mer (KH-52) precursor peptide containing the SIINFEKL epitope and a 46-mer (MP-46), and a 53-mer (KH-53) precursor peptide containing the MBP(79-87) epitope were synthesized (Supplementary Fig. 4). The 45-mer and the 46-mer precursor peptides correspond to the amino acid sequence of the β-globin exon region in which the SL8 or the MBP(79-87) epitopes are introduced in the construct used in Figure 1E. The 52-mer and the 53-mer precursor peptides correspond to the amino acid sequence of the β-globin intron region in which the SL8 and the MBP(79-87) epitope are introduced in the same construct used in Figure 1D. Importantly, preliminary studies demonstrated that the commercial enzymes used in these experiments were active and free of contaminant proteases, as shown by the ability of epoxomicin to completely inhibit the chymotrypsin- and trypsin-like activities of these preparations (Supplementary Table 1). KH-52 and KH-53 were then incubated *in vitro* with human constitutive 20S proteasome at 37°C for several hours, and the rates of peptide bond hydrolysis were assessed by measuring the generation of new amino groups with fluorescamine (35). Under these conditions, KH-52 and KH-53 were clearly degraded by h20Sc at linear rates over the entire time course of incubation (Fig. 5B). Moreover, the rates of peptide bond hydrolysis were not appreciably enhanced in the presence of REGγ and appeared to be only slightly lower than the rates measured when the proteasome associated with REGα/β (Fig. 5B, compare blue and red curves, left and right panels). Furthermore, indicative of absolute proteasome-dependent degradation, peptide hydrolysis was completely prevented in the presence of 20 μM epoxomicin (Fig. 5B, green curve, left and right panels). Similar results were also obtained when MP-45 and MP-46 precursor peptides were incubated under the same experimental conditions (Fig. 5C).

We then assessed the second hypothesis namely that REGγ stimulates 20S proteasome in a manner that leads to complete degradation of MHC class I epitopes. In fact, although binding of REGγ did not result in significantly increased rates of substrate hydrolysis (Fig. 5B and 5C), it may modify proteasome cleavage specificities to generate different patterns of peptide products as already extensively demonstrated for REGα/β (34). Such a change in enzymatic properties may potentially affect the formation of specific antigenic peptides such as SIINFEKL. To test directly this possibility, peptides released during KH-52 hydrolysis by equimolar amounts of 20S with and without REGγ were analyzed by tandem mass spectrometry (MS/MS). Importantly, shorter peptides (representing the great majority of proteasomal products (34,48,49)) were excluded to minimize false-positive identification and signals originating from small chemical compounds and therefore were not identified or quantified. In contrast, peptides with the correct size to bind to MHC-I heterodimers (i.e., 8-10 mers) were accurately analyzed. Using this approach, we identified 76 different peptides from KH-52, ranging in length from 8 to 16 residues and derived from the entire sequence of the substrate. Importantly, some peptides were generated exclusively by one form of the proteasome (i.e., 20S or REGγ-20S) (supplementary Table 2), while several others were released by both proteasomal forms. Although MS/MS does not provide quantitative information regarding the absolute abundance of the peptides detected, it is possible to assess their relative amounts by comparing the corresponding ion intensities measured in sequential MS/MS analyses. Therefore, we used ion intensities to quantify the relative amounts of single fragments generated from KH-52 in both degradation reactions. Remarkably, this analysis demonstrated that different amounts of peptide products are released by 20Sc and REGγ–20S (Fig. 5D). Most importantly, we demonstrated that the generation of SIINFEKL was strikingly suppressed in the presence of REGγ, as inferred by a two-log value reduction in its ion intensity (Fig. 5E).

All of these results unambiguously demonstrate that the regulator REGγ has the capacity to modify the cleavage properties of the 20S proteasome in the nuclear compartment in such a way that PTP-derived MHC class I antigenic peptides are destroyed rather than correctly processed and released.

### Knockout of REGγ gene causes tumor growth defect

Considering the previous results obtained with the human cell lines on the expression of REGγ, we decided to look at the REGγ mRNA level and REGγ expression in the mouse sarcoma MCA205 and mouse melanoma B16F10 cell lines and to compare them with those observed in the primary skin fibroblast cell line B6. We observed by qRT-PCR that REGγ mRNA was notably upregulated in MCA205 tumor cell line compared to the B6 cell line (Supplementary Fig. 5A). Furthermore, we noticed an overexpression at the protein level of REGγ in MCA205 murine cancer cell line compared to B6 cell line (Fig. 6A). To evaluate whether the extent of the difference in REGγ expression in murine cell lines contributes to antigen production and presentation, murine sarcoma and primary skin fibroblast cell lines were transiently induced to express the Globin-SL8-intron construct and were compared in terms of B3Z hybridoma activation. We observed that antigen presentation in sarcoma cancer cell line tested was lower than that in the fibroblast cell line supporting our previous results with the human cancer cell lines (Fig. 6A). To further depict the role of REGγ in tumorigenicity and cancer immune response, we genetically deleted REGγ in the MCA205 sarcoma cancer cell line (Fig. 6B, upper panel and Supplementary Fig. 5B). Then *Cas9*-REGγ MCA205 and MCA205 cells were transiently expressing the Globin-SL8-intron construct and were compared in terms of B3Z hybridoma activation. In this way we observed that *Cas9-*REGγ MCA205 clones induce a stronger activation of B3Z compared to the parental murine cell line (Fig. 6B, lower panel). In the same manner, and to support our above results showing that downregulation of REGγ induce a better cancer immune response due to the non-degradation of PTP-dependent antigens, rather than as a consequence of an overall increased expression of MHC-I molecules at the cell surface, we acid stripped cell surface class I molecules and measured the recovery of surface class I expression. The *Cas9-*REGγ MCA205 clone does not show any increase in MHC-I K^b^ molecules at the cell surface compare to the WT cancer cell line. Rather, a decrease in the amount of total MHC-I molecules is observed although these cells elicit a better cancer immune response compared to that obtained in presence of REGγ overexpression (Supplementary Fig. 5C). This result convincingly supports the idea that the effect of REGγ on antigen presentation takes place earlier than the loading of the antigenic peptides and the exporting steps of the MHC class I peptide complex.

**Figure 6:**
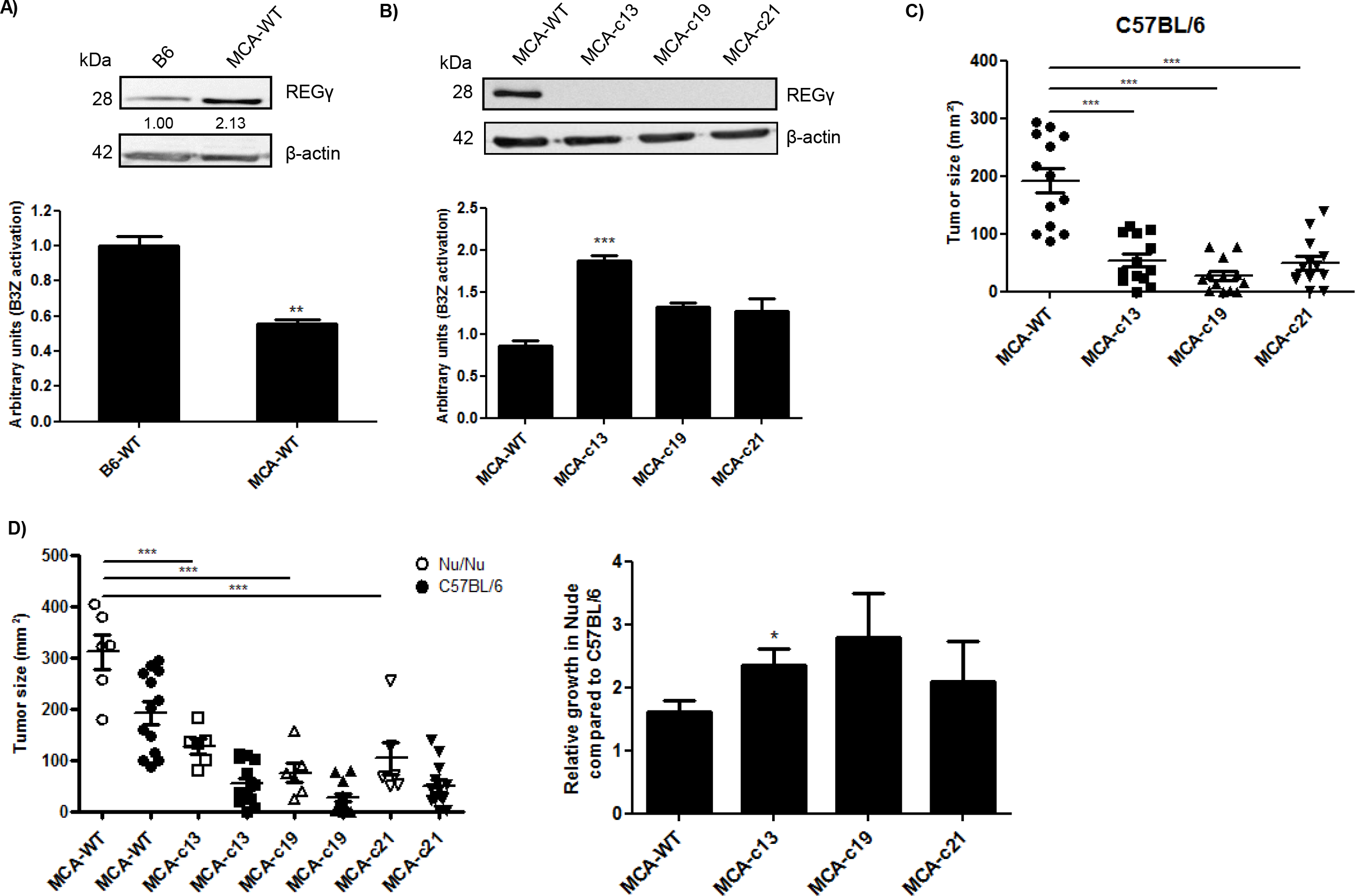
Knockout of REGγ gene causes tumor growth defect and changes on the tumor immunopeptidome. **a)** The B6 fibroblast and MCA205 sarcoma cell lines were transfected with β-Glob-intron-SL8 constructs. Cells were co-cultured with the SL8-specific CD8+ T cell hybridoma (B3Z) for 16 h. Upper panel shows the protein level of REGγ in the tumor cells assessed by Western Blot (quantification relative to the housekeeping protein β-actin). Lower panel shows the relative level of antigen presentation in the tumor cells. **b)** The MCA205 sarcoma cell line and the *Cas9-*REG◻ MCA205 clones were transfected with the β-Glob-intron-SL8 construct for 48 h. The cells were then co-cultured with the SL8-specific CD8+ T cell hybridoma (B3Z) for 16 h. Upper panel shows the protein level of REGγ in the tumor cells assessed by Western Blot (quantification relative to the housekeeping protein β-actin). Lower panel shows the relative level of antigen presentation in the tumor cells. **c)** Tumor size in area of sarcoma MCA205 WT and *Cas9-*REGγ MCA205 clones subcutaneously inoculated into the right flank of immunocompetent C57BL/6 mice. **d)** Tumor size in area of sarcoma MCA205 WT and *Cas9-*REGγ MCA205 clones subcutaneously inoculated into the right flank of immunodeficient Nu/Nu mice or immunocompetent C57BL/6 mice (left panel). Relative growth of MCA205 WT and *Cas9-*REGγ MCA205 clones in immunodeficient Nu/Nu mice compared to immunocompetent C57BL/6 mice. Data are given as mean ± SEM. *p<0.05, **p<0.01 (ANOVA with Tukey’s multiple comparaison test comparing all groups).

The proteasome regulator REGγ has been reported to have a specific effect on tumorigenicity. In fact it has been shown that REGγ contributes to the proliferation and metastasis of different cancer types. To gauge the effect of REGγ on cancer immune response and tumorigenicity *in vivo*, mice were subcutaneously injected into the flank with MCA205 WT cells or with several *Cas9-*REGγ MCA205 clones respectively. As shown in figure 6C, the *Cas9-*REGγ MCA205 clones originate smaller tumors than the parental cell line at day 20 after challenge. This result indicates that the loss of REGγ induces a decrease in the tumorigenicity of sarcoma cells. To validate that this decrease in tumorigenicity was due at least in part to a specific activation of the cancer immune response, as we observed *in vitro*, Nude/Nude mice were subcutaneously injected with MCA205 cells or several *Cas9-*REGγ MCA205 clones. As shown in figure 6D, the relative growth of the different *Cas9-*REGγ MCA205 clones in nude mice was higher than that of the WT cells. This difference in tumor growth between the *Cas9-*REGγ MCA205 clones and the parental cell line in immunodeficient nude mice demonstrates the crucial role of REGγ in tumorigenicity but also in antigen presentation and tumor immune responses. These *in vivo* results confirm all our previous *in vitro* finding clearly demonstrating that there is an inverse correlation between REGγ expression and antigen presentation in cancer and showing that cancer cells overexpress REGγ to destroy PTP-dependent antigenic peptides in order to reduce specific tumor immune responses.

## Discussion

This study describes for a first time a distinct and unexpected role of the REGγ-proteasome complex, which proved to be responsible for MHC-I antigens degradation in cancer. Specifically, we demonstrate an inverse correlation between REGγ expression and MHC class I antigen presentation in cancers, thus revealing a new function of this regulator in blunting immuno-responses by modulating the activity of the proteasome processing pathway *in vitro* and *in vivo*.

Contrary to our results, approximately twenty years ago, REGγ−/− mice were reported to exhibit neither positive nor negative effects in terms of antigen presentation. In fact, Barton et al. demonstrated that REGγ−/− mice did not show notable impairment in antigen presentation after viral γ infection (29). The only observed phenotypes reported so far for REGγ-/- mice so far are a small reduction in the numbers of specific CD8+ T lymphocytes and growth retardation (29,30). In this study, we clearly demonstrate for the first time a specific role for REGγ in MHC-I antigen processing and presentation. First of all, we show that REGγ acts as a negative regulator of MHC-I antigen presentation in cancer by destroying *in vitro* and *in cellulo* MHC-I peptides generated during proteasomal degradation of PTPs. Furthermore, we demonstrate that overexpression of REGγ negatively affects cancer immune response *in vivo*. Considering our results, it is therefore not surprising that Barton et al. did not observe any effects on antigen presentation in mice lacking REGγ. In fact, we observed a defect in MHC-I antigen presentation in cancer cell lines or in mice only when REGγ was naturally overexpressed. Indeed, when REGγ was knocked down in different cancer cell lines using either siRNA, miRNA-7 overexpression, cisplatin treatment, or CRISPR technology, we observed an important strengthening of CD8+ T cell activation and an enhanced immunological response against tumor progression *in vivo*. In contrast, in normal cell lines in which REGγ is not overexpressed or when we rescue the expression of REGγ in the A375 melanoma CRISPR cell line (A375cXI), we have been capable to inhibit MHC-I antigen presentation, at the same level observed in cancer cell lines naturally overexpressing REGγ. It is generally believed that only REGα/β in association with the immunoproteasome is able to affect the production of MHC-I epitopes. Strikingly, our results clearly demonstrate a close correlation between the expression of REGγ and tumor immune evasion. Thus, not only the heteroheptamer REGα/β plays a role in antigen presentation but REGγ does as well, although exerting an opposite effect. In fact, we proved that knocking out of REGγ reduces the growth of sarcoma MCA205 in immunocompetent mice. However, when we performed the same experiment in Nude nu/nu mice, which are deficient in T cells but not in B or innate immune cells, we did observe an increased tumor growth of several *Cas9-*REGγ MCA205 clones compared to the MCA205 WT cells that overexpress REGγ. This crucial experiment firmly established that knocking out REGγ induces the presentation of new PTP-MIPs that are recognized by CD8+ T cells, and so delaying the tumor growth. In fact, the tumor growth delay is likely to be a consequence of the escape from proteasomal destruction of new PTP-MIPs arising from translation of pre-spliced mRNAs leading to the apparition of epitopes derived from non-coding regions.

As mentioned above, many studies have reported that REGγ is overexpressed in several cancers (38–40). This overexpression constitutes a beneficial setting for tumors to proliferate and become metastatic. In fact, REGγ was recently shown to be involved in the degradation of some regulatory proteins. For example, the cyclin-dependent kinase inhibitor p21 is specifically degraded in an ATP- and ubiquitin-independent process via the REGγ-proteasomal pathway (18). Furthermore, two other cyclin-dependent kinases inhibitors, p16 and p14, are reportedly degraded via the same pathway (17). Even more importantly, REGγ has been shown to be involved in the MDM2-mediated p53 degradation process (20). These findings demonstrate the important role of REGγ in tumorigenicity by regulating cell proliferation and apoptosis. In this study, we demonstrate a new role for REGγ that is beneficial for cancer growth and development. Indeed, REGγ overexpression not only encourages cancer progression but, as confirmed by our data, is involved in reducing the amount of polypeptides that can serve as MHC-I epitopes, thus causing an inefficient immune response against tumor cell lines and finally resulting in the invisibility of cancer cells to the host immune system.

The REGα/β complex is largely located in the cytoplasm, while REGγ is mostly a nuclear proteasomal regulator (15). REGγ has been shown to be associated with the 20S proteasome in nuclear speckles (21). Moreover, many reports have demonstrated that the 20S proteasome and immunoproteasome are present in the nucleus and, more specifically, in clastosome structures (50). Considering that PTPs are produced by a nuclear translation event, it is clearly more beneficial for cells to process different PTPs in the same compartment where they are produced. In terms of the cancer hiddenness to the host immune system, it is also more advantageous for cancer cells to rapidly destroy tumor-associated PTPs (TA-PTPs) where they are produced rather than to give them the chance to be properly processed in the cytoplasm in a way that might generate class I epitopes and thus induce specific cancer immune responses. Therefore, by overexpressing REGγ in the nucleus where PTPs are located, cancer cells add another layer of regulation to the immune response. Indeed, we showed that by manipulating the localization of REGγ via overexpression of the ΔNLS variant of the regulator, we were able to restore a proper immune response against TA-PTPs. This identification of a specific site for TA-PTP degradation reinforced our recent hypothesis that, under normal conditions, MHC-I antigen epitopes are produced and processed in the nucleus, but under abnormal conditions such as cancer, TA-PTPs are still produced in the nucleus but are immediately processed by the REGγ-proteasomal complex in a specific way that minimizes the generation of MHC-I epitopes instead of be properly processed by different types of proteasomes such as the standard 20S proteasome, the immunoproteasome or a REGα/β-proteasomal complex, all of which were shown to be able to generate MHC-I epitopes (13,51–54)

The eukaryotic proteasome possesses three different peptidase activities. The β1 subunit exhibits caspase-like or peptidyl-glutamyl peptide hydrolyzing (PGPH) activity, the β2 subunit exhibits trypsin-like activity, and the β5 subunit exhibits chymotrypsin-like activity (55). When cells are treated with interferon gamma (IFNγ), the expression of three additional catalytic β-subunits is stimulated. All three active constitutive β-subunits are thus replaced, in newly assembled complexes, by a corresponding IFNγ induced β-subunits, specifically LMP2 (β1i), LMP7 (β5i) and LMP10 (β2i) (Mecl1), producing the so-called immunoproteasome (28). Although the heptameric REGα/β has been reported to stimulate all three proteasomal activities (12), the effect of REG γ on proteasomal cleaving specificities appear more complex and strictly related to the exact sequence of the fluorogenic short peptides used to assess them (27). The heptameric REGα/β regulator binds to 20S proteasome and immunoproteasome and in this way enhances generation of bind some, but not all, MHC-I epitopes. Two different explanations might be proposed to explain the ability of REGγ to decrease PTP-dependent antigen presentation: REGγ might (1) inhibit PTP processing via the proteasome complex or (2) enhance the destruction by 20S proteasome of MHC-I epitopes embedded in PTPs. Here, we propose that by binding to the nuclear 20S proteasome REGγ stimulates the degradation, rather than the proper processing, of MHC-I antigenic peptides by changing the peptidase activities of the 20S core particle. *In vitro* degradation assays, in fact, clearly showed that several polypeptides (ranging in length between 45 and 53 residues), with size and sequences corresponding to different PTPs, are efficiently hydrolyzed by REGγ-20S proteasomes at rates only slightly lower than those obtained with REGa/b-20S particles. These experiments, therefore, rule out the possibility that REGγ, by binding to 20S proteasome, may simply act as general inhibitor of proteasomal degradation of PTPs. In line with the alternative hypothesis that REGγ may induce destruction, rather than proper processing, of specific antigenic peptides, MS analysis demonstrated that 20S proteasome is able to efficiently process and release the immunodominant epitope SIINFEKL embedded in the precursor peptide KH-52, corresponding to the sequence of the β-globin intron region. Crucially, however, when the precursor peptide KH-52 was degraded by the 20S proteasome activated by REGγ, the generation of the immunodominant epitope SIINFEKL was completely abolished. Although our MS semiquantitative analysis precluded the recognition of small peptides generated following the fragmentation of SIINFEKL, the stimulation of proteasomal peptidase activities, resulting in the destruction of the antigenic peptide, appeared to be the more likely explanation for the inability to detect the immunodominant epitope in the presence of REGγ.

We recently reported that PTPs are a major source of MHC-I antigenic peptides for the endogenous and exogenous pathways. In fact, by inhibiting PTP degradation by the proteasome in tumor cell lines, we were able to increase the amounts of unprocessed PTPs entering the exogenous pathway, resulting in an increase in the activation of naïve CD8+ T cells (7,8). The role we unveiled of REGγ in the degradation of PTPs and specifically its ability to inhibit the endogenous MHC-I antigen presentation from TA-PTPs in tumors, leads to the hypothesis that this mechanism might also affect PTP-dependent exogenous MHC-I presentation. Indeed, by overexpressing REGγ in a CRISPR REGγ knock-out cell line, we showed that this cell line was no longer able to induce the proliferation of CD8+ T cells via the exogenous pathway. Moreover, mutated and ΔNLS-REGγ constructs played a less active role in PTP degradation, indicating that the main function of REGγ was to modulate the activity of the nuclear proteasomal complex, with strong inhibitory consequences not only on MHC-I endogenous antigen presentation but also on MHC-I exogenous pathway. Thus, we are driven to speculate that cancer cells might induce the overexpression of REGγ for both purposes: not only to inhibit MHC-I endogenous pathway but also to ensure that TA-PTPs have no chance to enter the MHC-I exogenous pathway and so to induce the proliferation of naïve CD8+ T cells.

In conclusion, the results of the present study, together with those of previous studies investigating REGγ functions, support the idea that tumor cell lines have an established system to positively regulate REGγ expression for specific purposes, such as to i) control cell cycle arrest, ii) control apoptosis, iii) control cell proliferation, and most importantly iv) inhibit endogenous and exogenous MHC-I antigen presentation by specifically degrading TA-PTPs encode MHC-I epitopes in order to become invisible to the host immune system.

## Acknowledgments

We thank Xiaotao Li (Department of Biochemistry and Molecular Biology, Institute of Biomedical Sciences, Shanghai, China) for the REGγ expression plasmids. The A375 and A549 cell lines were a gift from Dr. Fabrice André, Gustave Roussy Institute. The WM3526 and WM3682 cell lines were a gift from Levi Garraway. The HT29 and T84 colon cell lines were a gift from Dr. Fanny Jaulin from the Gustave Roussy Institute. We also thank Prof. Laurence Zitvogel for valuable comments on the manuscript. This work was funded by The Avenir program (INSERM), La Fondation ARC, and Fondation Gustave Roussy. BM is supported by the Fondation Gustave Roussy, and PA is supported by the Université Paris Sud.

