## supplemental Figures for "Tumors escape immunosurveillance by overexpressing the proteasome activator REGγ"

**Supplementary Figure 1**

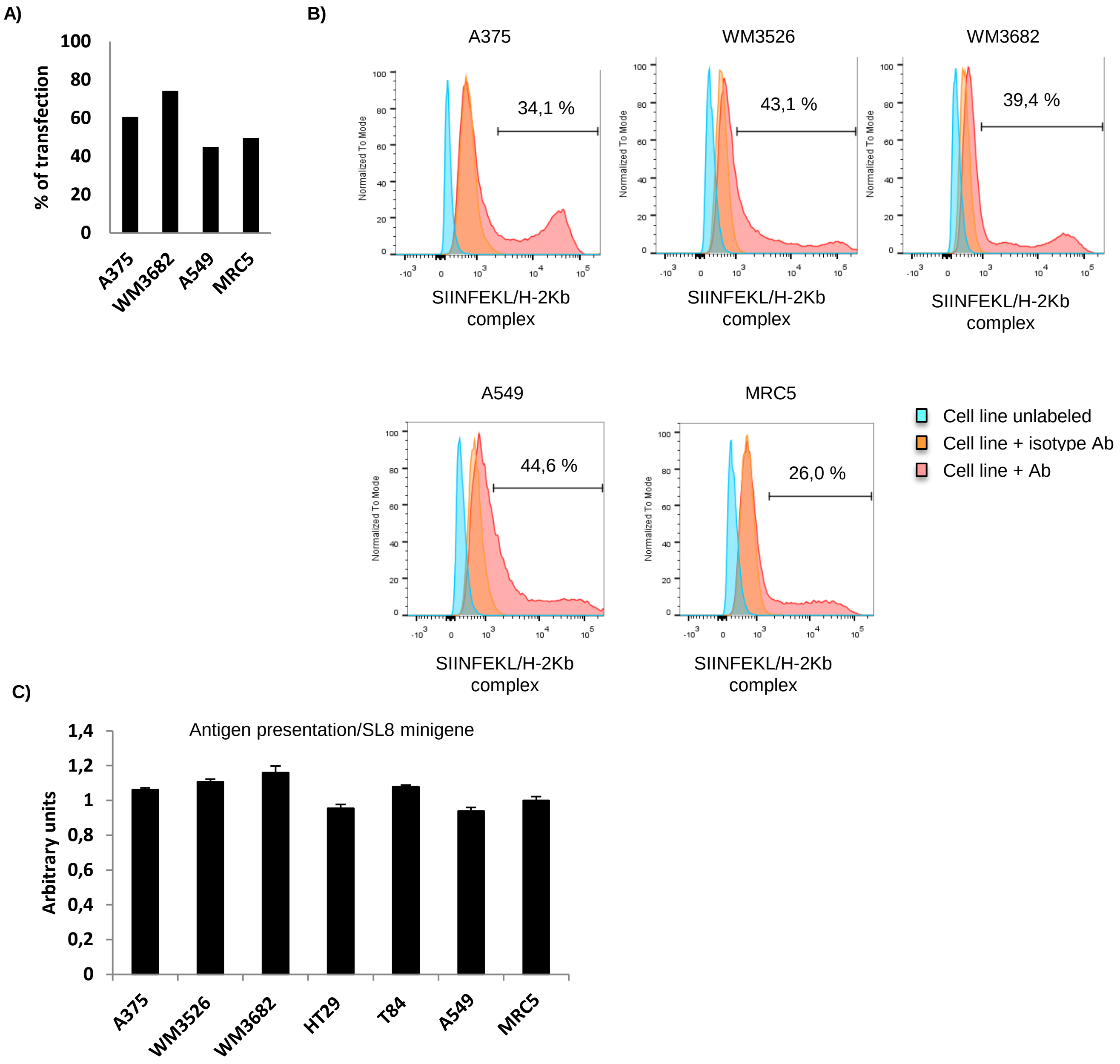

**Supplementary Figure 2**

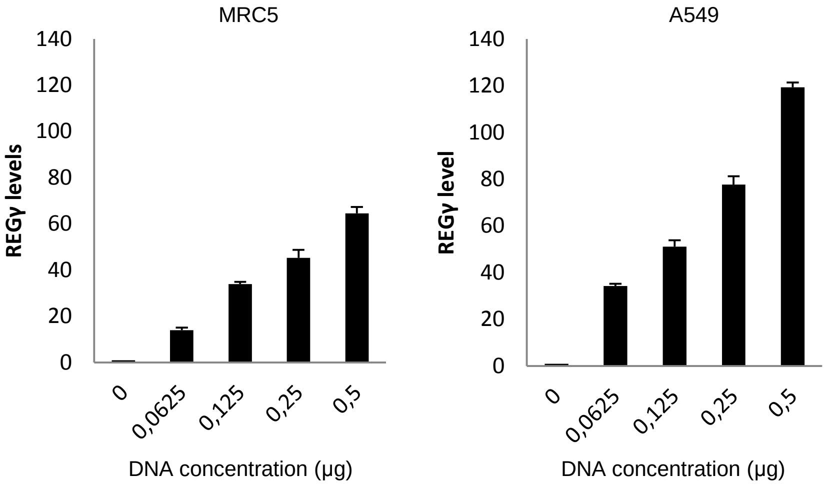

**Supplementary Figure 3**

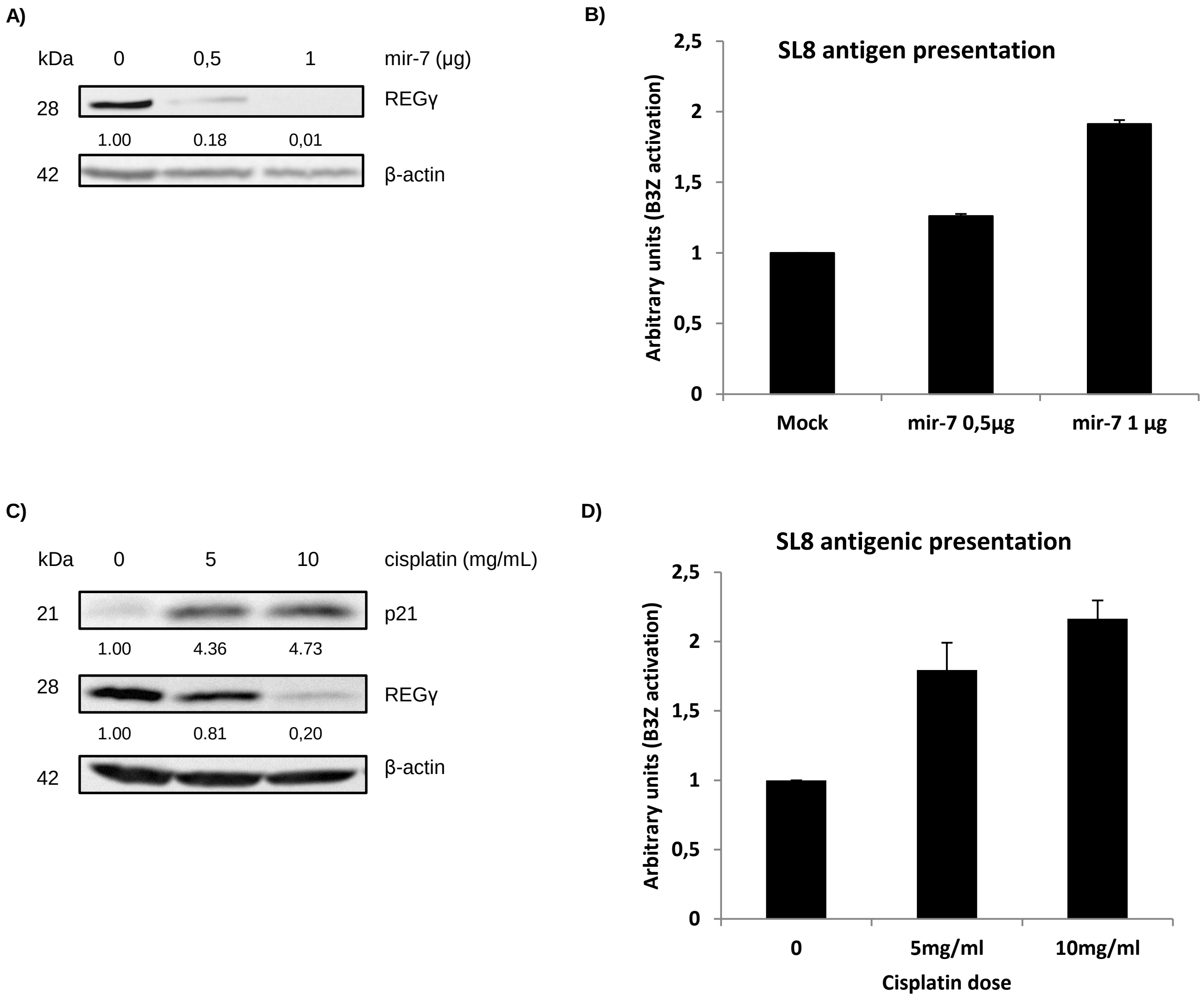

**Supplementary Figure 4**

**MP-45: Peptide exon-SIIN**
M V H L T D A E K S A V T **S I I N F E K L** A L W A K V N P D E V G G E A L G R L L V V Y P

**KH-52: Peptide intron-SIIN**
K V N V D E V G G E A L G R L V S R L Q D R **S I I N F E K L** F K E T N R N W A C G D R E D S W V S D R H

**MP-46: Peptide exon-MBP(79-87)**
M V H L T D A E K S A V T **D E N P V V H F F** A L W A K V N P D E V G G E A L G R L L V V Y P

**KH-53: Peptide intron-MBP(79-87)**
K V N V D E V G G E A L G R L V S R L Q D R **D E N P V V H F F** F K E T N R N W A C G D R E D S W V S D R H

**Supplementary Figure 5**

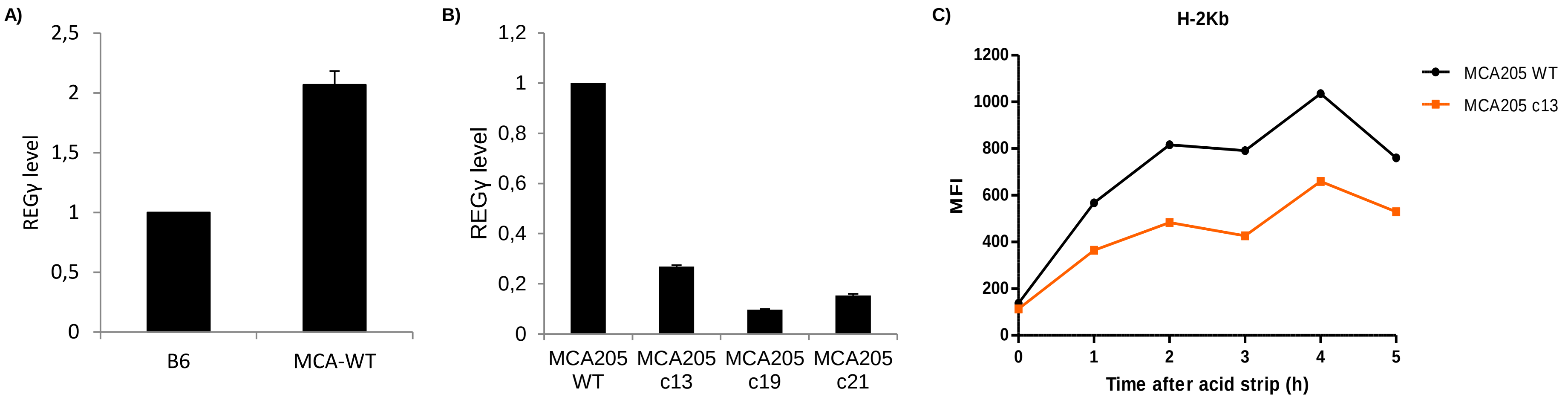
