## supplemental methods for "Tumors escape immunosurveillance by overexpressing the proteasome activator REGγ"

**Supplemental experimental procedures**

**Western blotting**

Cells were lysed in buffer containing 25 mM Tris–HCl, pH 7.6, 150 mM NaCl, 1% NP-40, 1% sodium deoxycholate and 0.1% SDS 20% and one complete protease inhibitor cocktail tablet (Thermo Scientific) per 10 mL of buffer. Cell lysates were centrifuged at 15,000 g for 10 min at 4°C. The protein contents in the supernatants were measured using a BCA Protein Assay kit (Pierce). Samples containing 40 μg of total protein were resolved in 15% SDS-PAGE gels. The proteins were transferred to Protan nitrocellulose membranes (Whatman) using a TRANS-BLOT Transfer Cell (Bio-Rad). Membranes were blocked in TBS-T containing 5% non-fat dried milk for 1 h at RT. Primary antibodies were diluted in 5% non-fat dried milk, and incubations were carried out at RT overnight. Primary antibodies were used at the following dilutions: REGγ, 1:1,000; p21, 1:1,000; REGα, 1:1,000; and actin (Sigma), 1:5,000. All other antibodies were obtained from Pierce Antibodies. HRP-conjugated secondary antibodies were incubated for 1 h at RT in 5% non-fat dried milk at a dilution of 1:2,500 for anti-rabbit and anti-mouse antibodies (Dako). Blots were developed using an enhanced chemiluminescence (ECL or West Dura (ThermoFisher)) Western blotting detection system (Ozyme).

**RNA preparation, RT and qPCR**

Total cellular RNA was extracted and purified using an RNeasy Mini kit (Qiagen) following the manufacturer’s instructions. RT was carried out with 0.5 μg of RNA using an iScript cDNA synthesis kit (Bio-Rad). A StepOne real-time PCR system (Applied BioSystems) was used for qPCR, and the reactions were performed with Power SYBR green PCR master mix (Applied BioSystems). The following human gene-specific primers were designed: PA28γ forward primer, 5’ – CACAAGTGAGGCAGAAGAC – 3’; PA28 γ reverse primer, 5’ –GAGATTCATGTCAGAGTGG – 3’; REGγ forward primer, 5’ – CATGTGGAGGACTATCGCCG – 3’; REGγ reverse primer, 5’ – CAGGATCATGTATGTAGAG – 3’; actin forward primer, 5’ –ATTGCCGACAGGATGCAGAA – 3’; actin reverse primer, 5’ –GCTGATCCACATCTGCTGGAA – 3’. The results were analyzed using StepOne software.

**Immunofluorescence**

Cells were plated at 2 × 10^5^ on 24 × 24 mm coverslips in 6-well plates. At 48 h post-transfection, the cells were washed briefly with 1× PBS-CM (0.1 mM **C**aCl_2_ and 1 mM MgCl_2_) and then fixed in 4% PFA for 30 min at RT, rinsed twice with 1× PBS-CM and permeabilized with PBS-CM 0.1% Triton X-100 for 5 min. Primary antibodies (see below) were incubated for 1 h at RT, and secondary antibodies were incubated for 30 min at RT, both in PBS-CM 10% FBS. The anti-REGγ antibody (Pierce Antibodies) and Flag tag antibody (a kind gift from Borek Vojtesek, Masaryk Memorial Cancer Institute, Brno, Czech Republic) were rabbit and mouse antibodies, respectively.

**DuoLink**

The A375cXI CRISPR cell line was plated at 2 × 10^5^ on 24 × 24 mm coverslips in 6-well plates. At 48 h post-transfection, DuoLink was performed using a DuoLink^®^ In Situ Detection Reagents FarRed kit (Sigma) following the instructions of the manufacturer. Primary antibodies were incubated in a humidity chamber for 1 h at RT: anti-Flag, 1/1,000^e^, and anti-20Sα4, 1/1,000^e^.

### Flow cytometry

Cell surface staining was performed on cultured human tumor cell lines that were previously transfected with a pcDNA3 plasmid containing the H-2K^b^ gene and stimulated with extracellular SIINFEKL synthetic peptide (Polypeptide, ovalbumin (257-264)) for 15 min at 37°C. Cells were stained with the anti-mouse OVA257-264 (SIINFEKL) peptide bound to H-2K^b^ APC (eBioscience) for 30 min. Controls were stained with mouse IgG1 K isotype control APC (eBioscience). DAPi was used for dead cell exclusion. Samples were acquired with a BD LSR II flow cytometer, and the results were analyzed using FlowJo software (version X.0.7).

### BMDC generation

BMDCs were generated from C57Bl/6 mice bone marrow precursors. Briefly, mouse femurs and tibia were removed, and the surrounding muscle tissue was cut and washed away in sterile PBS. Intact bones were placed in 70% ethanol for 2–3 min and then rinsed with sterile PBS. Epiphyses were removed with sterile scissors, and marrow was flushed with PBS/5% FBS. The suspension was pipetted several times to disperse clusters of cells, and the cells were centrifuged at 1200 rpm for 5 min. Cells were then seeded at 1 × 10^7^ cells/mL in 145-mm Petri dishes for 6 days in IMDM (Sigma) supplemented with 10% heat-inactivated FBS, 0.5% penicillin/streptomycin, 1% L-glutamine, and β-mercaptoethanol (50 nM, Life Technology) and enriched with 50 ng/mL J558 supernatant. After 3 days of culture, the medium was replaced.

### BMDC cross-presentation

On day 6 of BMDC differentiation, 0.3 × 10^6^ BMDCs were placed in contact with human tumor cells transfected with the plasmid pcDNA3 containing the β-globin intron gene, empty plasmid or with tumor cells stimulated with the SIINFEKL synthetic peptide. As a positive control, OVA protein was added to the BMDCs. As a negative control, BSA was added to the BMDCs. After 24 h, the BMDCs were washed with medium and cultured with 0.3 × 10^6^ B3Z cells overnight in a 24-well plate. B3Z activation was quantified as previously described in the T cell assay methods.

**Immunoprecipitation**

Cells were washed in 1× ice-cold PBS and lysed in buffer containing 0.5% NP-40, 150 mM NaCl, 20 mM Tris-HCl pH 7.4 and 5% glycerol. Cell lysates were centrifuged at 15,000 g for 10 min at 4°C. Magnetic beads were washed 3 times in lysis buffer and incubated for 1 h at 4°C under gentle agitation with cell supernatant and normal serum of the same species of IP antibody. The supernatant was then incubated for 1 h at 4°C under rotation with the primary antibody targeting REGγ, which was used at 1:100 (Pierce Antibodies). Magnetic beads were added for overnight incubation at 4°C. Finally, the beads were washed 3 times in lysis buffer. The samples were separated on a Western blot gel to evaluate protein precipitation.
