## Supplemental table 1 for "Tumors escape immunosurveillance by overexpressing the proteasome activator REGγ"

|  | V_max_ (nmol/mg*min) | | |
| --- | --- | --- | --- |
| Substrate | h20Sc | h20Sc  + epoxomicin | h20Sc  + REGγ |
| Suc-LLVY-amc | 297.7 | 0.0 | N.D. |
| Bz-VGR-amc | 269.5 | 0.0 | 387.9 |
| **Supplementary Table 1. Chymotrypsin-like and trypsin-like specific activities of the human constitutive 20S proteasome (h20Sc).** Specific activities of the h20Sc and h20Sc proteasomes activated by REGγ were determined as described in the Materials and Methods. Epoxomicin was used at final concentrations of 2 µM and 20 µM to assess chymotrypsin-like and trypsin-like activities, respectively. | | | |
