## Supplemental figure legends for "Tumors escape immunosurveillance by overexpressing the proteasome activator REGγ"

**Supplementary Figure 1: Inverse correlation between the expression of REGγ and antigen presentation in multiple cancer cell lines. a)** A375, WM3682, A549 and MRC5 cells were transfected with the Glob-intron-SL8 construct vector for 48 h. The transfection efficacy was analyzed by FACS. **b)** FACS analysis of H-2K^b^ expression at cancer cell surface. Human tumor cell lines were transfected with a construct expressing the H-2K^b^ gene, stimulated with extracellular SIINFEKL synthetic peptide for 15 min and processed for FACS staining. Controls were stained with mouse IgG1 K isotype control APC. **c)** All cell types were transfected *in vitro* with a SL8 minigene-expressing construct. The cells were incubated with the B3Z T cell hybridoma for 16 h. The data show the average of at least three independent experiments ± SD minus the values from mock-transfected cells. Free SL8 peptide was added to the cells to ensure that T cell assays were performed under non-saturated conditions and that the expression of MHC class I molecules was not affected.

**Supplementary Figure 2: Exogenous REGγ overexpression decreases MHC class I antigen presentation.** MRC5 (left panel) and A549 (right panel) cell lines were transfected with a REGγ WT-expressing construct (0.0625 µg, 0.125 µg, 0.25 µg and 0.5 µg). REGγ mRNA levels were analyzed by qPCR. REGγ mRNA levels were normalized to β-actin mRNA levels. Experiments were performed in triplicates. Data are expressed as the mean ± SEM from three technical replicates.

**Supplementary Figure 3: Knockdown and knock-out of the regulator REGγ promotes antigen presentation. a)** Western blotting was performed to analyze and quantify (relative to the housekeeping protein β-actin) REGγ protein levels in the A375 cell line. The cells were transfected with different concentrations of a miR-7 expression construct (0.5, 1 and 2 μg) for 48h. Protein levels are indicated below each gel. **b**) A375 cells were transfected with a construct expressing Glob-intron-SL8 (0.5 μg) for 48 h. The cells were incubated with the B3Z T cell hybridoma for 16 h. The data show the average of at least three independent experiments ± SD minus the values from mock-transfected cells. **c)** Western blot analysis and quantification (relative to the housekeeping protein β-actin) of REGγ and p21 protein levels were assessed in the A375 cell line. Cells were treated with different concentrations of cisplatin for 16 h. As reported previously, REGγ protein levels declined after cisplatin treatment. **d)** A375 cells were transfected with a Glob-intron-SL8 construct (0.5 μg) for 48 h. At 36 h post-transfection, A375 cells were treated overnight with cisplatin at different concentrations. The cells were then incubated with the B3Z T cell hybridoma for 16 h. The data show the average of at least three independent experiments ± SD minus the values from mock-transfected cells.

**Supplementary Figure 4: REGγ promotes the degradation of MHC class I PTP-derived antigenic epitopes.**

Amino acid sequences of the different polypeptides used for *in vitro* degradation, with SIINFEKL and MBP(79-87) epitopes indicated in red in the middle of each precursor peptide.

**Supplementary Figure 5**: **Knockout of REGγ gene causes tumor growth defect and changes on the tumor immunopeptidome.**

**a)** REGγ mRNA levels were analyzed by qPCR in sarcoma MCA205 and the fibroblast B6 cell lines and normalized to β-actin mRNA levels. Experiments were performed in triplicate. Data are expressed as the mean ± SEM from three technical replicates. **b)** REGγ mRNA levels were analyzed by qPCR in sarcoma MCA205 and the different Cas9-REGγ MCA205 clones and normalized to β-actin mRNA levels. The fibroblast B6 cell line was used as a reference. Experiments were performed in triplicate. Data are expressed as the mean ± SEM from three technical replicates. **c)** Kinetics of the recovery of H-2Kb molecules at the cell surface of MCA205 WT and Cas9-REGγ MCA205 clone after acid strip followed by flow cytometry with anti- H-2Kb antibody. MFI, Mean Fluorescent Unit.
