## Supplementary figures and images for "Tumors escape immunosurveillance by overexpressing the proteasome activator REGγ"

### Supplemental table 1

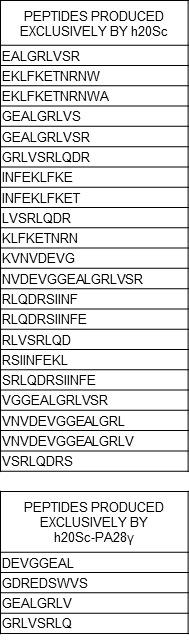

**Supplementary Table 2.** List of peptides produced exclusively by h20Sc or h20Sc-REGγ
